# Multiset partial least squares with rank order of groups for integrating multi-omics data

**DOI:** 10.1101/2022.08.30.505949

**Authors:** Hiroyuki Yamamoto

## Abstract

As more multi-omics data such as metabolome, proteome and transcriptome data are acquired, statistical analyses to integrate multi-omics data has been required. Multivariate analysis for integration of multi-omics data has been proposed such as multiset partial least squares (PLS). However, when we want to extract scores about the rank order of groups such as severity of disease as well as difference of groups, we could not always extract scores with rank order of groups by using multiset PLS. Therefore, we proposed multiset PLS with rank order of groups (multiset PLS-ROG) to integrate multiple omics dataset. We visualized multiple omics data by using multiset PLS-ROG in two studies of multi-omics and multi-organ derived metabolome data. After we visualize data and focus on a specific score with phenotype of interest, it is important to select compounds to identify biomarker candidates or make biological inferences. In ordinary principal component analysis (PCA) or PLS, we could select statistical significantly compounds correlated with PC or PLS score by using PC or PLS loadings. However, multiset PLS loading has not been used because its statistical property has not been clarified. Then, we clarified statistical property of multiset PLS and multiset PLS-ROG loading, and we could identify statistically significant compounds by using statistical hypothesis testing. We implemented multiset PLS-ROG to R loadings package, freely available in CRAN.

## 1. Introduction

Recently, many omics studies such as transcriptome, proteome and metabolome and integrated analysis for these multi-omics data have been reported [1, 2]. In an early study of multi-omics, Ishii et al [3] visualized multi-omics data by mapping gene expression, proteome, and metabolome data of *E. coli* onto a single metabolic pathway. This approach has extended to visualize relationships between each omics layer by connecting the regulatory relationships as trans-omics [4]. Many other multi-omics studies such as integration of metagenome and metabolome for gut microbiota [5] have been actively reported. Multi-omics studies were also not only applied for making biological inference in experimental biology, but also for genome-wide association between genome and protein or metabolite such as quantitative trait in clinical research [6].

For statistical analysis of omics data, various univariate analysis methods such as *t*-test and multivariate analysis methods have been applied. However, a standard statistical method for integrating multi-omics data has not yet been established, compared to the routine application of certain statistical analyses such as hierarchical clustering in transcriptomics and principal component analysis (PCA) in metabolomics. In the context of the necessity for standard statistical analysis methods for integrating multi-omics data, multiset multivariate analysis has been applied since the early 2010 to integrate multi-omics data [7]. One of the most representative tools of multiset multivariate analysis for integrating multi-omics data is mixOmics [8] of the Bioconductor R package that has been cited by above one thousand papers in 2022. The mixOmics is one of the standard statistical analysis tools for integrating multi-omics data.

In mixOmics package, multi-set partial least squares discriminant analysis (PLS-DA) is mainly adopted. While variable importance in projection (VIP) has often been used to select important compounds (i.e., genes, proteins and metabolites) in PLS-DA, correlation coefficient between multiset PLS-DA scores and each compound levels is used to select important compounds in multiset PLS-DA of mixOmics package. On the other hands, PC or PLS loadings have been also used to select important metabolites in metabolomics. We previously clarified the statistical properties of the PC or PLS loadings and applied statistical hypothesis testing of PC [9] or PLS loading [10] to select statistically significant metabolites. As well as PCA or PLS, the statistical properties of multiset PLS loadings are expected to be clarified because we can select statistically significant compounds by using statistical hypothesis testing of multiset PLS loading.

In practical data analysis, we have sometimes an additional information about groups such as the severity of the disease or the concentration of the drug to be administered. When we applied PLS to this type of data, the rank order of the groups is not necessarily represented in the PLS scores. Therefore, we previously proposed PLS with rank order of groups (PLS-ROG). In this study, we extended PLS-ROG to multiset PLS-ROG for integrating multiple omics dataset. We not only organized multiset PLS-ROG from a theoretical perspective but also applied multiset PLS-ROG to multi-omics dataset. As a practical application of multiset PLS-ROG, we reanalyzed the multi-omics data of proteome and metabolome data from human serum samples on severity of COVID-19 and reanalyzed the multiple metabolome data from rabbit samples derived from different organs and plasma on the effects of drugs on hyperlipidemia. From practical data analysis in two studies, we confirmed the effects of the parameters on multiset PLS-ROG scores and on the change of highly ranked statistically significant compounds. Multiset PLS-ROG also has the advantage that is easy to implement, and can be computed using loadings package of R, which is freely available from the CRAN website.

## 2. Methods

### 2-1. Multiset PLS-ROG

PLS is formulated as maximization of covariance between score for explanatory variable (i.e., omics data) and for response variable (i.e., group information) [10,11]. And PLS-ROG [10] is formulated as well as PLS, but differs from PLS in that a smoothing term for the group-wise mean of the scores for response variable is added to the constraint condition of PLS. Multiset PLS is also formulated as maximization of sum of covariance between scores for all combinations of each explanatory variables and between score for response and each explanatory variable. As well as multiset PLS, multiset PLS-ROG is formulated as maximization of sum of covariance and almost the same constraint condition as PLS-ROG as follows,

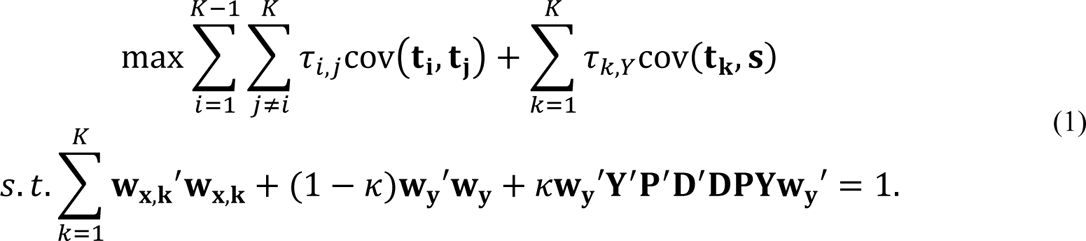

*K* is the number of datasets. The parameter *τ_i,j_* represents the strength of the connection between i-th and j-th omics dataset and *τ_k,Y_* represents the strength of the connection between k-th dataset and the response variable. These parameters *τ_i,j_* and *τ_k,Y_* are set so that these values sum up to 1 as follows,

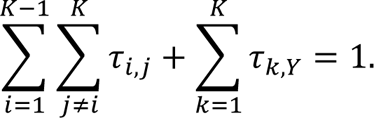

In the implementation, we summarized *τ_i,j_* and *τ_k,Y_* as the following matrix **τ**,

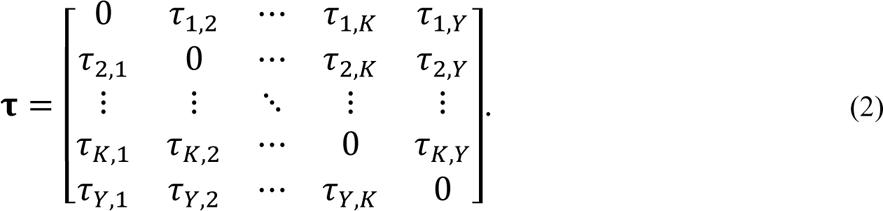

The **τ** is a symmetric matrix whose diagonal elements are zero because the same dataset is not connected to each other. The cov(**t_k_**, **s**) means the covariance between the k-th score for explanatory variable **t_k_**=**X_k_w_x,k_** and the score for response variable **s=Yw_y_**. The **w_x,k_** is the weight vector for the k-th data matrix **X_k_** and **w_y_** is the weight vector for the dummy matrix corresponds to the response variable. The k-th data matrix **X_k_** includes samples in rows and metabolites in column, and the matrix **Y** includes samples in row and group number in column. For example, when the number of groups is g, the matrix **Y** is set as follows,

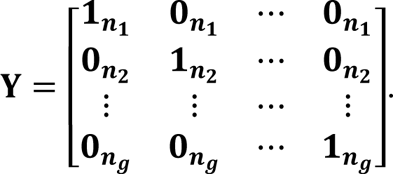

The elements of the matrix **Y** are vectors, for example, **1_*n*_1__** means an n_1_-dimensional vector with 1 as an element and **0_*n*_1__** means an n_1_-dimensional vector with 0 as an element. In the practical calculations, **X** and **Y** were mean-centered matrix, and the matrix **X** was further divided by the standard deviation for each metabolite levels (i.e., autoscaling) in this study. The first order differential matrix **D** and averaging matrix **P** are set for the g-class classification problems as follows,

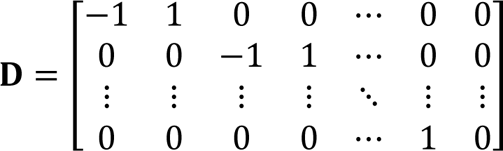

And

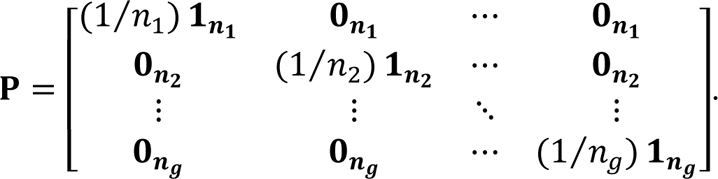

*κ* is a smoothing parameter with 0 ≤ *κ* < 1. When κ is closer to 1, the rank order of groups is more likely to appear in the multiset PLS-ROG scores because the effect of smoothing term of *k***W_y_′Y′P′D′DPYW_y_′** is relatively large in eq. (1). When *κ* equals to 0, multiset PLS-ROG equals to the multiset PLS.

The maximization problem under constraint condition in eq. (1) can be written by using the method of Lagrange multiplier as follows,

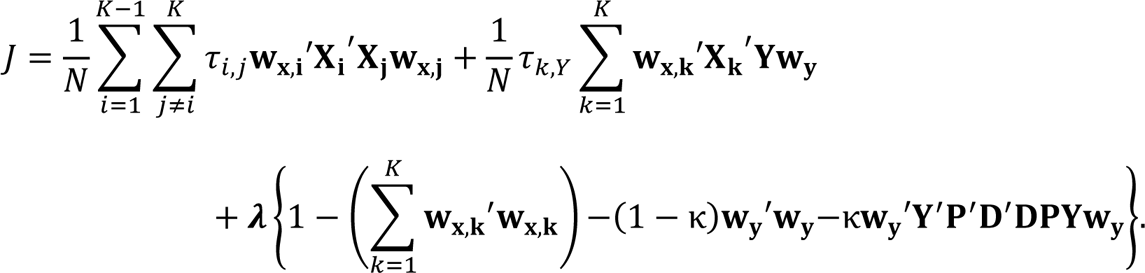

This equation is partially differentiated by the weight vector of **w_x,1,_ w_x,2, …,_ w_x,K_** and **w_y_** respectively, and setting it to 0.

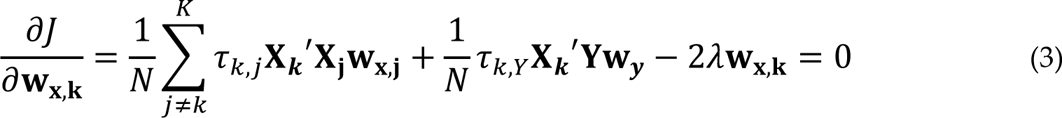

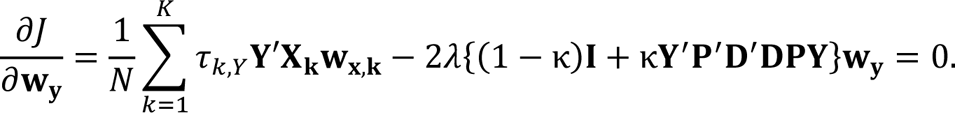

where k=1, 2, …, K and **I** is the identity matrix. Here, we review the meaning of λ. We multiplied by **w_y_** from the left of the lower expression in eq. (3),

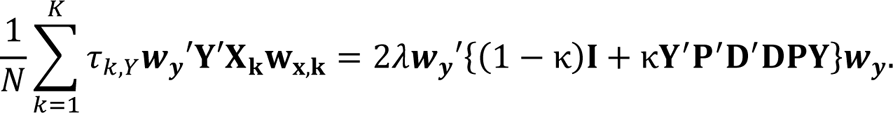

The left side of the above equation is a weighted sum of covariance between the score for explanatory **t_k_** and response variable **s**, and λ can be written as follows,

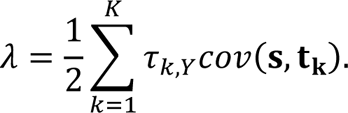

Finally, the eq. (3) can be written as the following generalized eigenvalue problem,

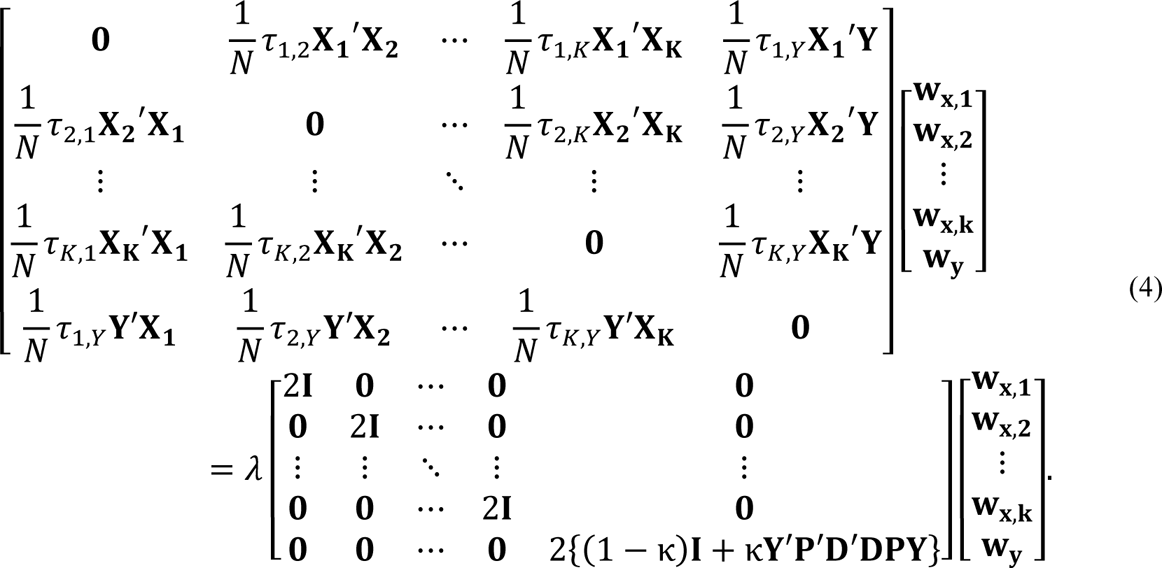

We focus on the component with the positively larger eigenvalue. For example, the component corresponds to the largest positive eigenvalue is the first component, and the component corresponds to the next largest positive eigenvalue is the second component.

The corresponding eigenvector is the weight vector **w** for each component. And, the multiset PLS-ROG score for the k-th dataset and the score for the response variable can be calculated with **t_k_**=**X_k_w_x,k_** and **s=Yw_y_**.

So far, we have considered the case where we connected between explanatory variables for each dataset, and between the response variable and explanatory variable in each dataset (Fig.1(a)). We also consider the case where we don’t connect between explanatory variables for each dataset (Fig.1 (b)). If the connection between the explanatory variables for each datasets were not made, *τ_i,j_* equals to 0 in equation (1). This eliminates the term 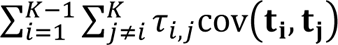 in eq (1) and can be written as follows,

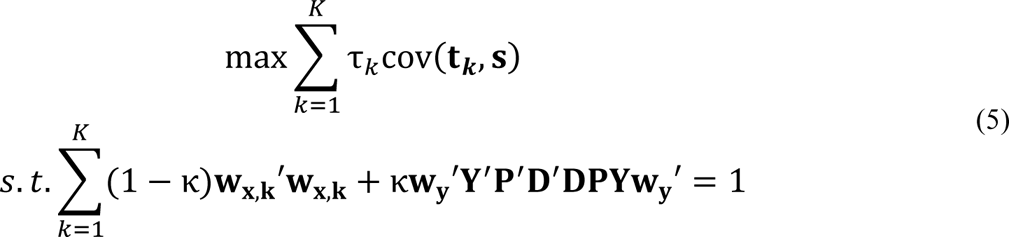

**Figure 1.**
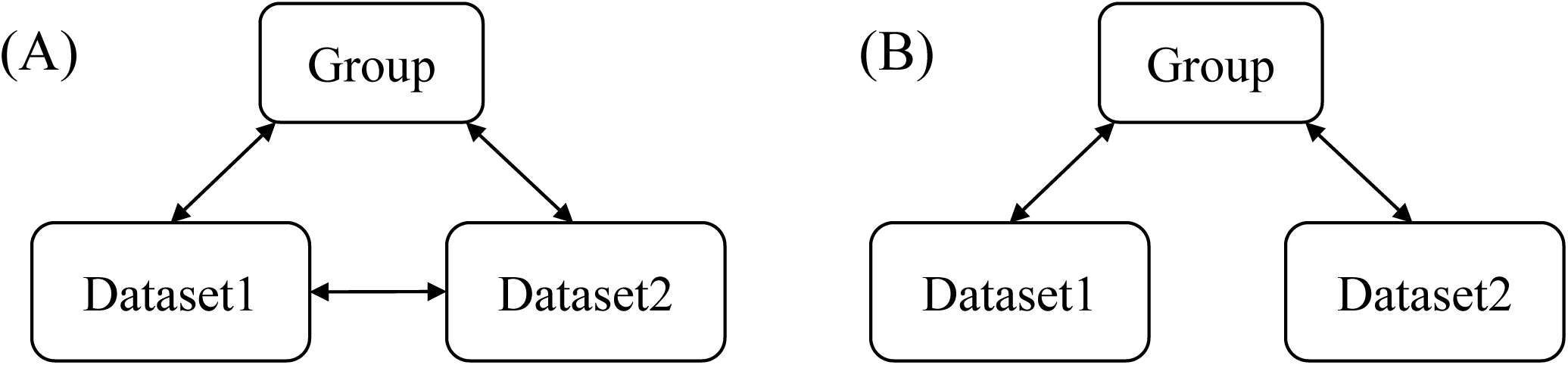
Diagram of connection between dataset and group information. (A) Connections between each omics datasets, and between each omics dataset and group information (B) Connections between each omics dataset and group information only.

When we set *τ_k_* as 1, the summation of covariance in eq. (5) can be re-written as covariance of the summation as follows,

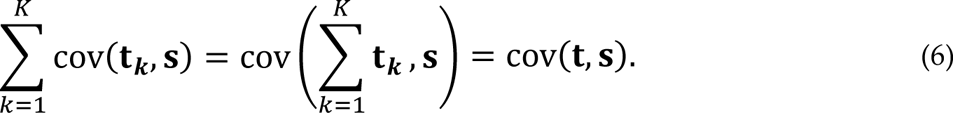

In the eq. (6), covariance between sum of the scores **t_k_** for the explanatory variables in each dataset k and the score for response variable is equivalent to the covariance between the score **t**, combining each dataset in the column direction into a single data matrix, and the score for response variable. In this case, we can compute multiset PLS-ROG by using PLS-ROG for this combined data matrix when samples in each row and compounds in each column. Multiset PLS-ROG score for explanatory variables in each dataset k can be computed as **t_k_**=**X_k_w_x,k_** by using the divided weight vector **w_x,k_** and each dataset **X_k_** (k=1,2,…,K).

In this study, we developed multiset PLS-ROG in supervised case and explained the computational framework for multiple datasets. Additionally, multiset PLS in unsupervised case, that connected between omics datasets only, can be calculated in the same framework.

### 2-2. Statistical property of multiset PLS-ROG loading

We visualize omics data with multiset PLS-ROG score and focus on that score associated with the phenotypes such as group information, and then select important compounds corresponding to this score. We can select relative important compounds by using the multiset PLS-ROG weight vector **w_x_** in eq (4). However, we cannot set a specific threshold for multiset PLS-ROG weight and select important compounds by using a statistical criterion because multiset PLS-ROG weight is set for computational convenience but not statistically meaning. In PCA and PLS, we clarified statistical properties of PC weight or PLS weight and were able to select statistically significant compounds by using statistical hypothesis testing of PC or PLS loading. As with PCA and PLS, it is important to clarify the statistical properties of the multiset PLS-ROG weight in order to select significant compounds by using statistical criteria.

To clarify the statistical property of multiset PLS-ROG loading, we first consider the weighted sum of covariance C*_x,k,p_* between the scores for the explanatory and response variables and the p-th metabolite in the k-th dataset.

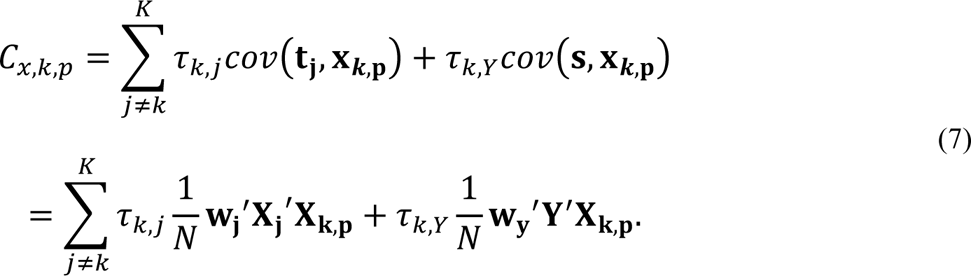

We plugged in the equation, 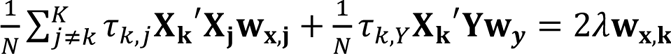, that is the transposed equation of a part of the eq. (3), into the eq. (7). Finally, the eq. (7) can be written as follows,

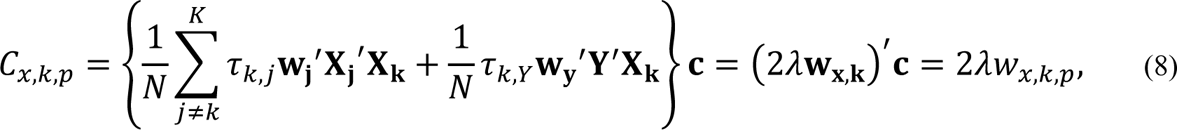

where **c** is the column vector in which the *p*-th element is 1 and the others are 0. From the eq. (8), we were able to confirm that multiset PLS-ROG weight **W_x.k_** is proportional to the weighted sum of correlation coefficient between the score for explanatory variable **t_j_** and *p*-th variable in k-th dataset **x_k,p_**, and between response variable **s** and **x_k,p._** And using the eq. (6), the weighted sum of covariance C*_x,k,p_* can be written as follows,

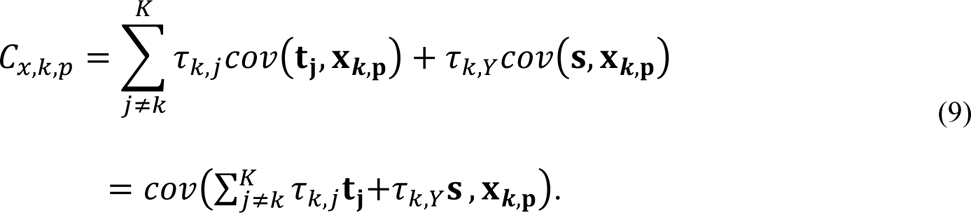

Fron the eq. (9), the weighted sum of covariance C*_x,k,p_* equals to the covariance between the weighted sum of score for explanatory **t** and response variable **s**. Based on these results, we defined the multiset PLS-ROG loading *R_x,k,p_* as follows,

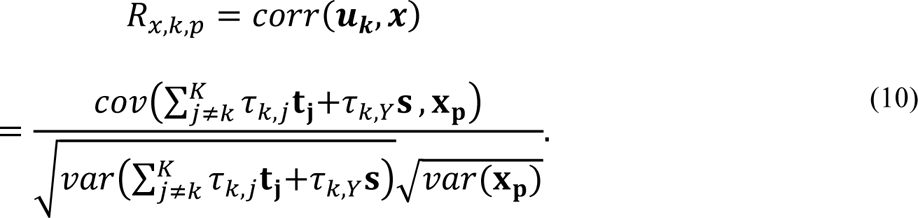

When each compound levels **x_k,p_** of p-th compound in k-th dataset are centered and scaled to unit variance (i.e., autoscaling), the var(**x_k,p_**), variance of **x_k,p_**, is 1. And, the variance of the weighted sum of scores can be regarded as a constant because the weighted sum of scores does not depend on the p-th compound of x_k,p._ Therefore, the relationship between the multiset PLS-ROG loading *R_x,k,p_* and the weighted vector **w_x,k_** also can be written as follows,

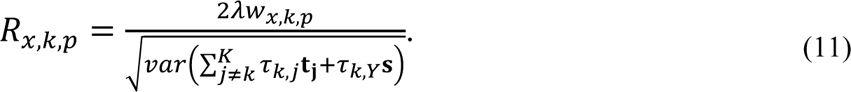

In our previous study of PCA, PLS and PLS-ROG, *p*-values of loadings can be calculated by using *t*-distribution because these loadings can be defined as correlation coefficient itself. As well as these approaches, the statistical hypothesis testing of factor loading in the multiset PLS and PLS-ROG can be performed by using that 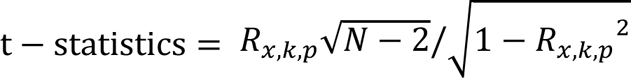 follows a *t*-distribution.

### 2-3. Implementation and computation

All statistical analysis in this study was computed by using R ver. 4.3.3, and we implemented the multiset PLS and multiset PLS-ROG to R loadings package. The loadings package is freely available in CRAN web site [https://cran.r-project.org/web/packages/loadings/]. We computed generalized eigenvalue problem by using geigen package. The *q*-value of multiset PLS-ROG loading was computed by Benjamini-Hochberg method.

## 3. Results

### 3-1. Multi-omics data in COVID-19 study

We re-analyzed multi-omics data of proteome and metabolome in COVID-19 study by Shen et. al. [11] as an application of multiset PLS-ROG. This dataset includes three groups of healthy control, mild and severe COVD-19. Our goal is to integrate and visualize multi-omics data and find score associated with severity of COVID-19, and then find compounds (proteins or metabolites) associated with the severity of COVID-19. We performed multiset PLS-ROG with the parameter κ as 0.999, default parameter of PLS-ROG in loadings packages, and with the weight of connectivity between group and each datasets are set to the same value as τ_1,2_=t_1,Y_=t_2,Y_=1/3.

Multiset PLS-ROG score for explanatory variables in the proteome (Fig.2 (A_1_)) and metabolome (Fig.2 (B_1_)) datasets showed that the specific samples of healthy subjects (JBDZ2, JBDZ3, and JBDZ6) located at a distance from many other samples in first multiset PLS-ROG score, and these three samples might be considered as outliers. We also confirmed three outlier samples in first and second score by unsupervised multiset PLS (Fig.S1) and supervised multiset PLS (Fig.S2). These results suggest that there is a strong association between proteome and metabolome data for specific samples. We also could confirm the rank order of groups, i.e., the severity of COVID-19, in first (Fig.2 (C_2_)) and second (Fig.2 (C_3_)) multiset PLS-ROG score for response variable. However, group differences and rank order of groups were not found in first multiset PLS-ROG scores of explanatory variables and rank order of groups was found slightly in second multiset PLS-ROG score of explanatory variables, but not clearly. This trend showed similar in multiset PLS scores (Fig S2).

**Figure 2.**
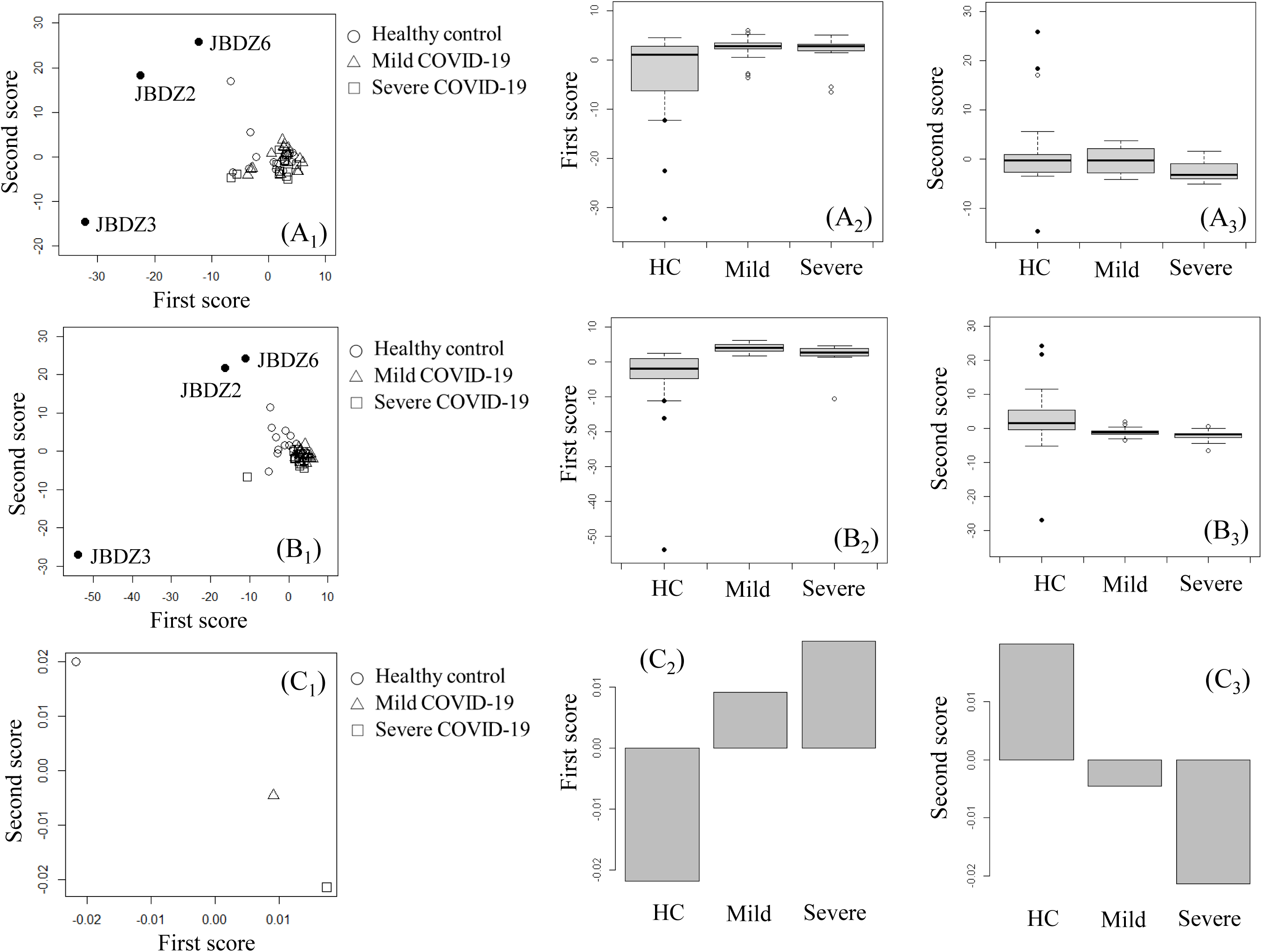
First and second multiset PLS-ROG scores in COVID-19 study. Three outlier samples of JBDZ2, JBDZ3 and JBDZ6 were blacked out. (A_1_) A scatter plot of first and second multiset PLS-ROG score for explanatory variables in proteome dataset. (◯) Healthy control (HC), (△) mild COVID-19, (□) severe COVID-19. A box and whisker plot of (A_2_) first multiset PLS-ROG score and (A_3_) second multiset PLS-ROG score for explanatory variables to HC, mild and severe COVID-19 groups in proteome dataset. (B_1_) A scatter plot of first and second multiset PLS-ROG score for explanatory variables in metabolome dataset. A box and whisker plot of (B_2_) first multiset PLS-ROG score and (B_3_) second multiset PLS-ROG score for explanatory variables to HC, mild and severe COVID-19 groups in metabolome dataset. (C_1_) A scatter plot of first and second multiset PLS-ROG score for response variables. A bar plot of (C_2_) first multiset PLS-ROG score and (C_3_) second multiset PLS-ROG score for response variable to HC, mild and severe COVID-19 groups.

One possible reason why group differences and rank order of groups do not appear clearly in multiset PLS scores of explanatory variables is that the relationship between groups and each omics data is weak. Then, we confirmed the correlation coefficient between first multiset PLS-ROG score for response variable and for explanatory variable. The correlation coefficient between first multiset PLS-ROG scores for proteome and metabolome data is 0.835, and the correlation coefficient between group and proteome data is 0.35 and the correlation coefficient between group and metabolome data is 0.427, respectively when τ_1,2_=t_1,Y_=t_2,Y_=1/3. This result showed that the correlation between multiset PLS-ROG score for response and explanatory variables was small, and as a result, the rank order of groups was not shown in multiset PLS-ROG score for explanatory variables. Moreover, we showed the change of correlation coefficient between group and each omics dataset when the weight of connectivity τ_1,Y_ and τ_2,Y_ was changed simultaneously from 0.3 to 0.5 in Fig.3. This result showed that the higher the weight of the connectivity between group and each omics dataset, the higher the correlation coefficient between the first multiset PLS-ROG score for response variable and for explanatory variable, and the gradually lower the correlation coefficient between the first multiset PLS-ROG scores of proteome and metabolome data for explanatory variable.

**Figure 3.**
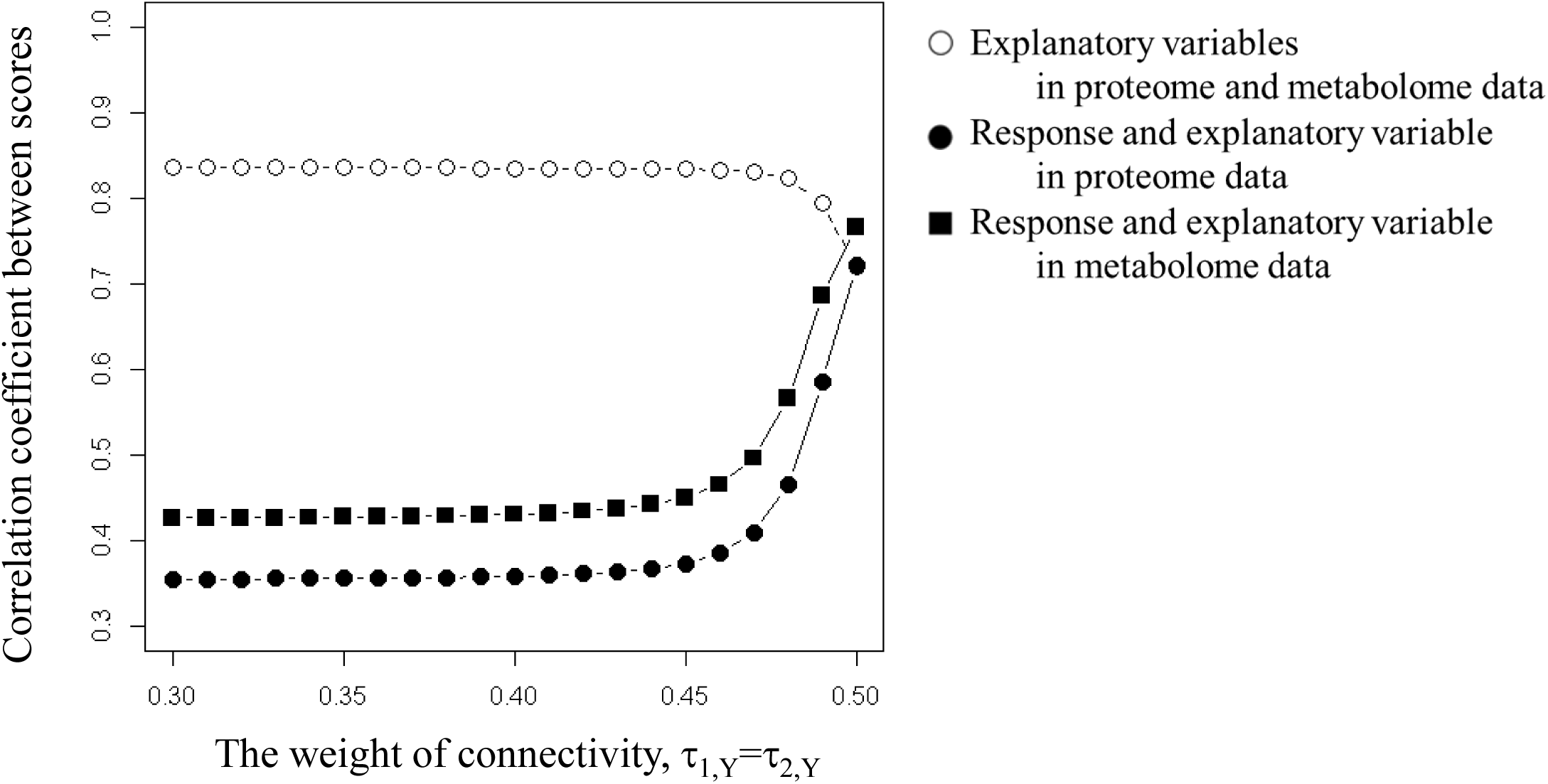
Correlation coefficient between multiset PLS-ROG score for response and explanatory variables when τ_1,Y_=τ_2,Y_ are set from 0.3 to 0.5 in 0.01 increments. (◯) Correlation coefficient between first multiset PLS-ROG score for explanatory variables in proteome and metabolome data, (●) correlation coefficient between first multiset PLS-ROG score for response and explanatory variable in proteome data (■) and correlation coefficient between first multiset PLS-ROG score for response and explanatory variable in metabolome data.

The results so far suggest that not only smoothing parameter κ but also the weight of connectivity τ greatly affect to score in multiset PLS-ROG. Then, we performed multiset PLS-ROG when the weight of connectivity between group and each omics datasets was set as τ_1,Y_=t_2,Y_=0.48 (Fig.S3), 0.49 (Fig.S4) or 0.50 (Fig.S5) in which the correlation coefficient changed significantly (Fig.3). In Fig.4, we showed the first multiset PLS-ROG score foxr explanatory variable of proteome data at τ_1,Y_=t_2,Y_=0.48, 0.49 or 0.50 to clearly represent how the score changes with the parameter τ. This result showed that the effect of outliers becomes smaller and the rank order of the groups becomes clearer in first multiset PLS-ROG score as the parameter τ_1,Y_ and τ_2,Y_ was increased from 0.48 to 0.50. And we picked up the top 3 highly correlated proteins with multiset PLS-ROG score by using multiset PLS-ROG loading. The first, second and third rank proteins were P12111 (R=-0.7966, p=7.645 × 10^-12^, q=6.048 × 10^-9^), P13727 (R=-0.7250, p=3.839 × 10^-9^, q=1.504×10^-6^) and P24821 (R=-0.7195, p=5.704×10^-9^, q=1.504×10^-6^) at τ=0.48, and were Q14520 (R=0.6648, p=1.889×10^-7^, q=1.494×10^-4^), Q15113 (R=-0.6374, p=8.422 × 10^-7^, q=2.350 × 10^-4^) and P12111 (R=-0.6363, p=8.913 × 10^-7^, q=2.350 × 10^-4^) at τ=0.49 and were Q14520 (R=0.6731, p=1.167×10^-7^, q=9.233×10^-5^), P12259 (R=0.6589, p=2.636×10^-7^, q=1.042×10^-4^) and Q15113 (R=-0.6118, p=3.007×10^-6^, q=7.929×10^-4^) at τ=0.50.

**Figure 4.**
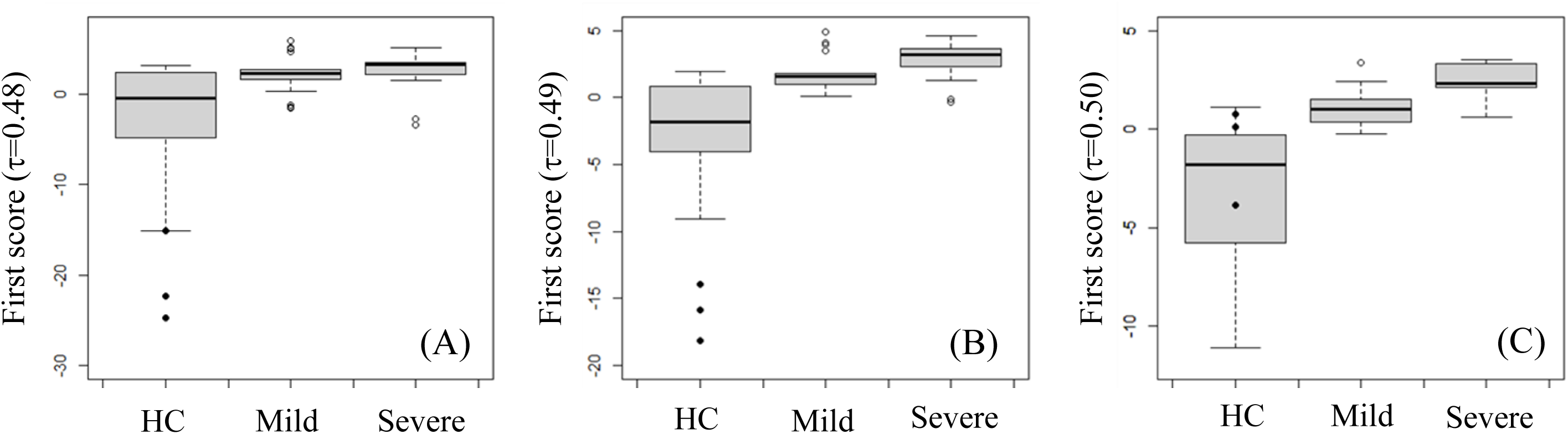
A box and whisker plot of first multiset PLS-ROG score for explanatory variables to HC, mild and severe COVID-19 groups at (A) τ_1,Y_=τ_2,Y_=0.48, (B) 0.49 and (C) 0.50 in proteome dataset.

We confirmed outliers in the top 3 ranked proteins of P12111 (Fig5(A)), P13727 (Fig,5(B)) and P24821 (Fig.5(C)) at τ_1,Y_=t_2,Y_=0.48 but not confirmed rank order of groups clearly. On the other hands, we confirmed rank order of groups in the top 3 ranked proteins of Q14520 (Fig.5(D)), P12259 (Fig.5(E)) and Q15113 (Fig.5(F)) at τ_1,Y_=t_2,Y_=0.50 clearly, but outliers were less impressive than top 3 proteins. We could confirm both rank order of groups and (although small impression) outliers in P14520 and Q15113. P14520 and Q15113 ranked top 1 and 2 at τ_1,Y_=t_2,Y_=0.49, which was between τ_1,Y_=t_2,Y_=0.48 where the effect of outliers was significant, and τ_1,Y_=t_2,Y_=0.50 where the effect of rank order groups was significant. These results suggested that, as well as the effect of outliers becomes smaller and the rank order of the groups becomes clearer in first multiset PLS-ROG score as the parameter τ_1,Y_ and τ_2,Y_ was increased from 0.48 to 0.50, we could also confirm same tendency in highly ranked proteins identified by multiset PLS-ROG loading as the parameter τ_1,Y_ and τ_2,Y_ was increased from 0.48 to 0.50. In proteome data, the top 3 with the highest correlation proteins in first multiset PLS-ROG loading at τ=0.50 were Q14520 of hyaluronan-binding protein 2 (Fig.5(D)), P12259 of coagulation factor V (Fig.5(F)), and Q15113 of Procollagen C-endopeptidase enhancer 1 (Fig.5(E)). The original paper for this multi-omics data of COVID-19 study discussed the proteins Q14520 and P12259, but not the protein Q15113. One reason that these proteins were not focused on might be that the levels of two proteins of Q14520 and P12259 increased but the level of the protein Q14126 decreased with the severity of COVID-19. In metabolome data, while we could confirm group difference between healthy control and COVID-19 in first multiset PLS-ROG score (τ_1,Y_=t_2,Y_=0.50) for explanatory variables in metabolomics, we could not confirm the rank order of groups clearly (Fig.S5 (B_2_)). The highest correlated metabolite in first multiset PLS-ROG loading at τ_1,Y_=t_2,Y_=0.50 was 5-methyluridine (R=-0.7360, p=1.694×10^-9^, q=1.580×10^-6^), also known as ribothymidine (Fig.S6). As well as the first multiset PLS-ROG score for explanatory variables, we could confirm the clear difference between healthy control and COVID-19 groups in 5-methyluridine but could not confirm the difference between mild and severe COVID-19, that was, could not confirmed the rank order of groups related with the severity of COVID-19. In the original paper of COVID-19 study [11], 5-methyluridine was not discussed. We inferred the reason was that this metabolite, like protein Q15113, increased but not decreased with the severity of COVID-19.

**Figure 5.**
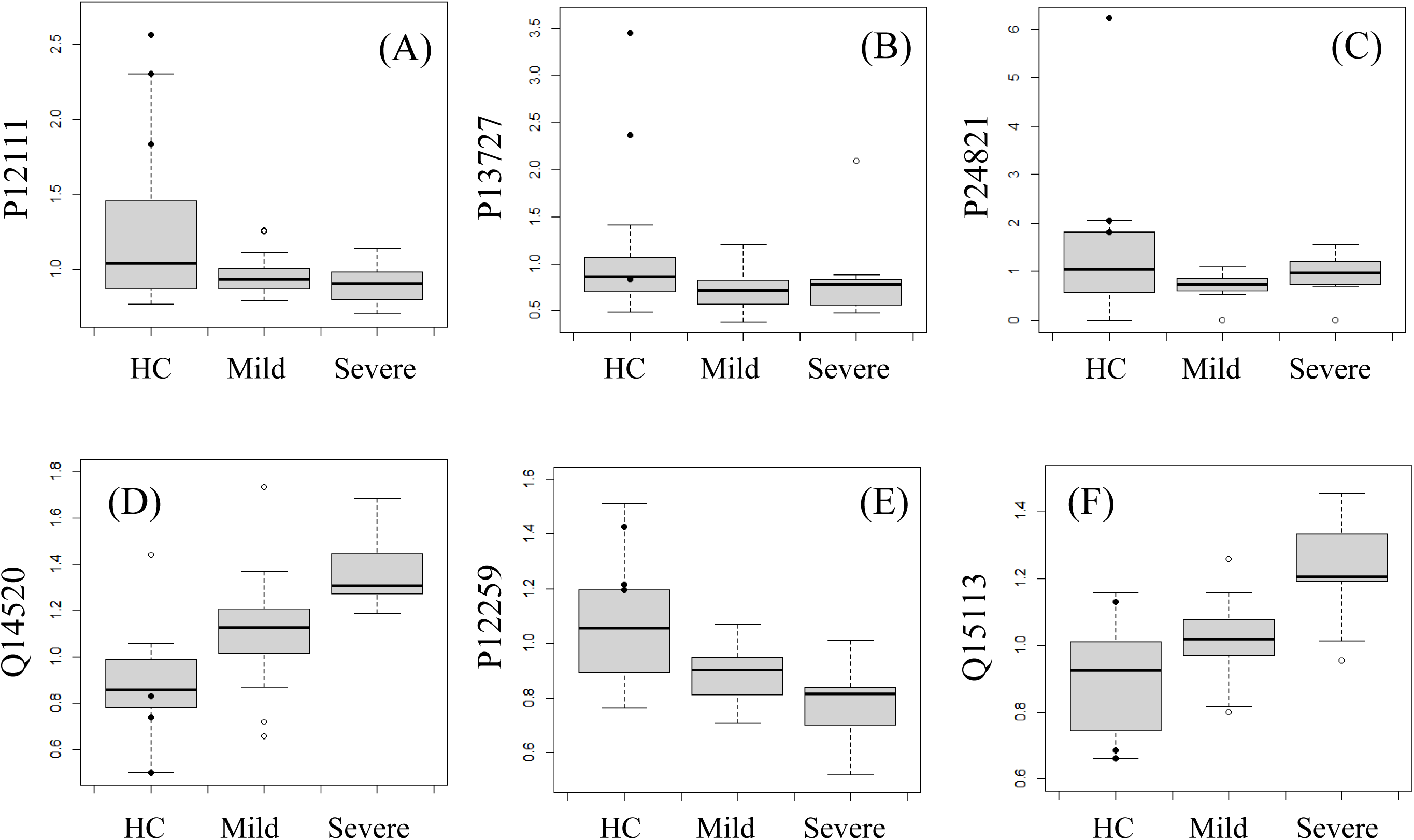
A box and whisker plot of top 3 ranked proteins, (A) P12111, (B) P13727, (C) P24821, (D) Q14520, (E) P12259 and (F) Q15113 by using multiset PLS-ROG loading at τ_1,Y_=τ_2,Y_=0.48, 0.49 and 0.50.

### 3-2. Multi-organ metabolome data in hyperlipidemia study

We re-analyzed multi-organ metabolome data of rabbit plasma, liver, heart muscle and brain samples in hyperlipidemia study [13]. This dataset has three groups of wild type, hyperlipidemia rabbit and hyperlipidemia rabbit with drug administration, with three samples in each group. The main purpose of this hyperlipidemia study is to visualize the effects of drugs on various organs and plasma in hyperlipidemic rabbits. Specifically, we want to find multiset PLS-ROG score that show the pattern that the hyperlipidemia group is approaching to wild type by drug administration, i.e. the pattern showing rank order of wild type, hyperlipidemia with drug administration and hyperlipidemia groups. After we find the multiset PLS-ROG score showing desirable pattern of rank order of groups, we can identify metabolites associated with drug effects by the corresponding multiset PLS-ROG loadings.

As in the previous section of COVID-19 study, we first performed multiset PLS-ROG with the parameter κ as 0.999 and all weight of connectivity was set to the same value, τ_i,j_=t_i,Y_=t_j,Y_ = 0.1 where i=1,⋯,4 and j=1,⋯,4 and i≠j (Fig. 6). We could confirm the group difference between hyperlipidemia with drug administration and other groups of wild type and hyperlipidemia group in first multiset PLS-ROG and multiset PLS score (Fig.S7), and could confirm rank order of groups in second multiset PLS-ROG and multiset PLS score of a part of explanatory, liver and heart muscle, and response variables. However, we could not confirm rank order of groups in first PLS-ROG score.

**Figure 6.**
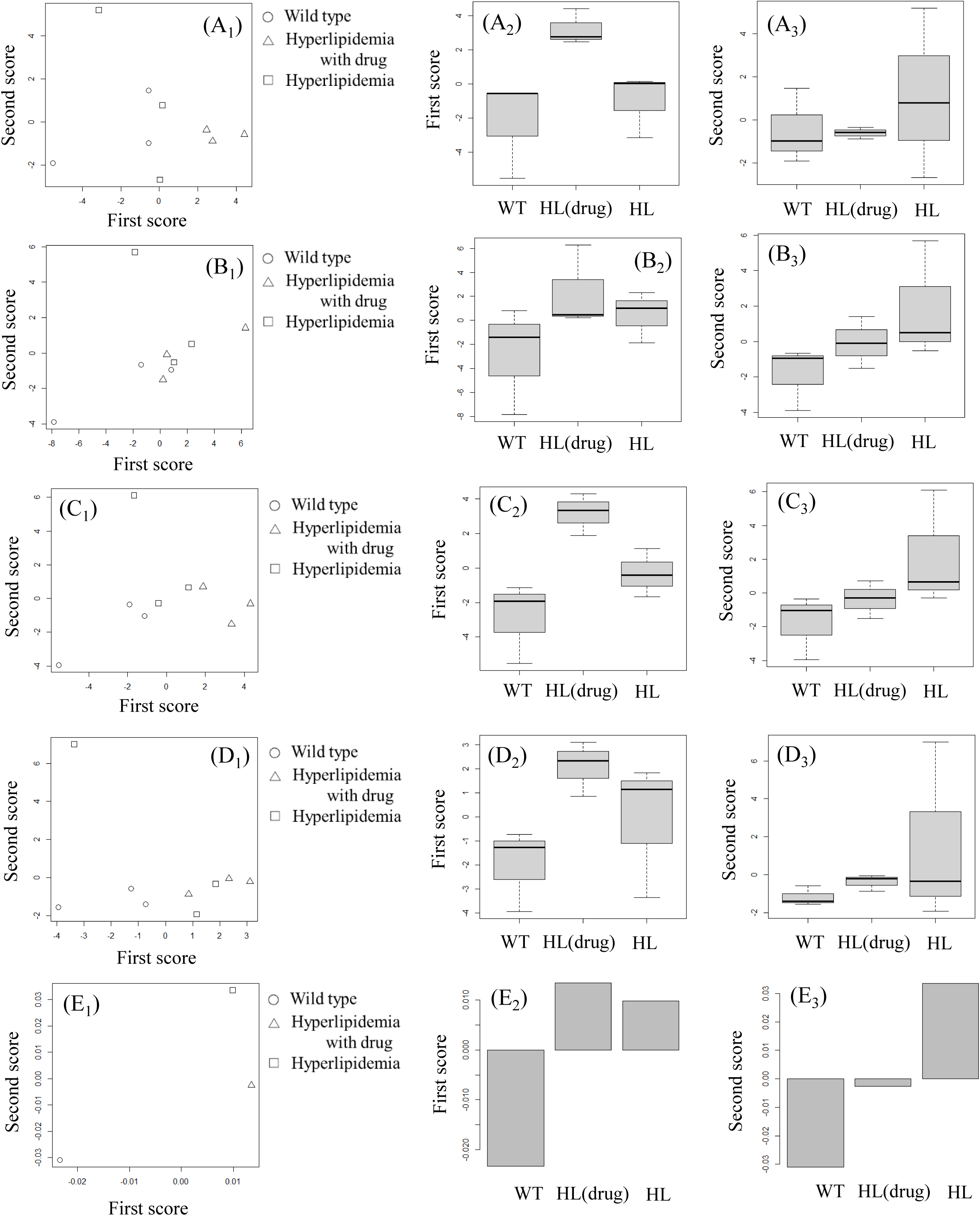
First and second multiset PLS-ROG scores (τ = 0.1) in hyperlipidemia study. (◯) Wild type (WT), (△) hyperlipidemia (HL) rabbit administered with drug, (□) HL rabbit. (A_1_) A scatter plot of first and second multiset PLS-ROG score for explanatory variables in plasma samples. A box and whisker plot of (A_2_) first multiset PLS-ROG score and (A_3_) second multiset PLS-ROG score for explanatory variables to WT, HL administered with drug and HL in plasma samples. (B_1_) A scatter plot of first and second multiset PLS-ROG score for explanatory variables to WT, HL administered with drug and HL in liver samples. A box and whisker plot of (B_2_) first multiset PLS-ROG score (B_3_) second multiset PLS-ROG score for explanatory variables to WT, HL administered with drug and HL in liver samples. (C_1_) A scatter plot of first and second multiset PLS-ROG score for explanatory variables in heart muscle samples. A box and whisker plot of (C_2_) first multiset PLS-ROG score and (C_3_) second multiset PLS-ROG score for explanatory variables to WT, HL administered with drug and HL in heart muscle samples. (D_1_) A scatter plot of first and second multiset PLS-ROG score for explanatory variables in brain samples. A box and whisker plot of (D_2_) first multiset PLS-ROG score and (D_3_) second multiset PLS-ROG score for explanatory variables to WT, HL administered with drug and HL in brain samples. (E_1_) A scatter plot of first and second multiset PLS-ROG score for response variables. A box and whisker plot of (E_2_) first multiset PLS-ROG score and (E_3_) second multiset PLS-ROG score for response variable to WT, HL administered with drug and HL.

As with the previous section, to clarify the effect of the weight of connectivity to multiset PLS-ROG score, we showed the change in correlation coefficient between group and each omics dataset when the weight of connectivity τ_i,Y_ and τ_j,Y_, hereinafter denoted as τ, was changed simultaneously from 0.1 to 0.25 (Fig.7). In this hyperlipidemia study, we used the mean value of the correlation coefficients between the multiset PLS-ROG scores for explanatory variables because a specific organ was connected to multiple other organs, e.g., plasma was connected to liver, heart muscle and brain. In Fig.7, we could confirm that the weight of connectivity affects greatly to the correlation coefficient between multiset PLS-ROG scores for response and explanatory variables as well as the previous section in COVID-19 study. We could also confirm that the correlation coefficients between second PLS-ROG score for explanatory variables of liver and heart muscle and for response variable were slightly decreased at τ=0.23 in Fig.7 (B). This depends on the fact that the pattern of the rank order of groups of wild type, hyperlipidemia with drug administration and hyperlipidemia groups was swapped from second PLS-ROG score to first PLS-ROG score from τ=0.22 to 0.23. In Fig.7 (B), we also confirmed that the correlation coefficient between second PLS-ROG score for explanatory variables of liver and heart muscle and for response variable are high and similar tendency with a variation of the parameter τ. Because statin is known to inhibit cholesterol production in the liver and have an effect on myocardial infarction, it is reasonable that the effect of statin is more clearly confirmed in the liver and heart muscle. To confirm the impact of parameter τ on the multiset PLS-ROG in more detail, we performed multiset PLS-ROG at τ =0.23 (Fig. S9), 0.24 (Fig. S10) and 0.25 (Fig. S11). First multiset PLS-ROG scores at τ=0.23, 0.24 and 0.25 for response and explanatory variables (Fig.8) were shown to clarify the effect on the score with respect to a change in the weight of connectivity. As the parameter τ increased from 0.23 to 0.25, we could confirm the rank order of groups of wild type, hyperlipidemia with drug administration and hyperlipidemia groups in first multiset PLS-ROG score for liver, heart muscle and brain clearly, and we could not confirm the rank order of groups but confirm pattens that hyperlipidemia group was approaching to the hyperlipidemia with drug administration group for plasma. As described above, statin have been known to effect on the liver and muscles, and also known to be effective not only in myocardium but also in brain infarction [14].

**Figure 7.**
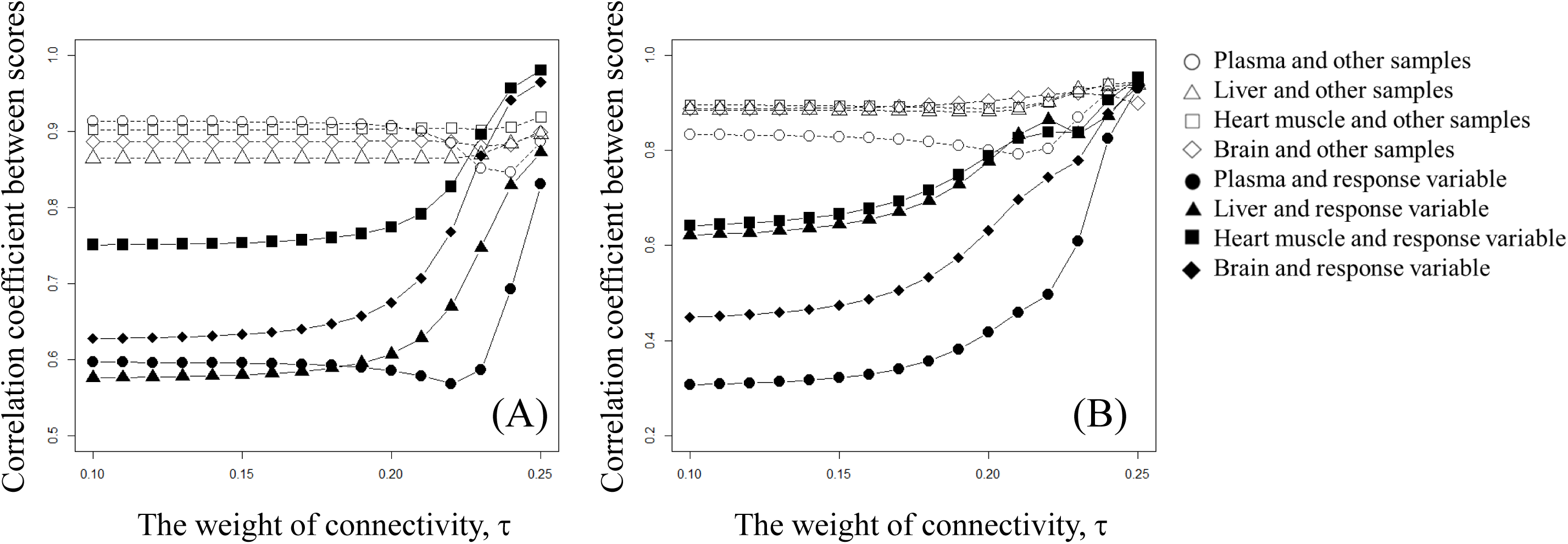
Correlation coefficient between multiset PLS-ROG score for response and explanatory variables when τ is set from 0.1 to 0.25 in 0.01 increments. Mean of correlation coefficient between first multiset PLS-ROG score for explanatory variables in (◯) plasma, (△) liver, (□) heart muscle, (⃟) brain and other samples and correlation coefficient between first multiset PLS-ROG score for response and explanatory variable for (●) plasma, (▲) liver, (■) heart muscle and (♦) brain samples.

**Figure 8.**
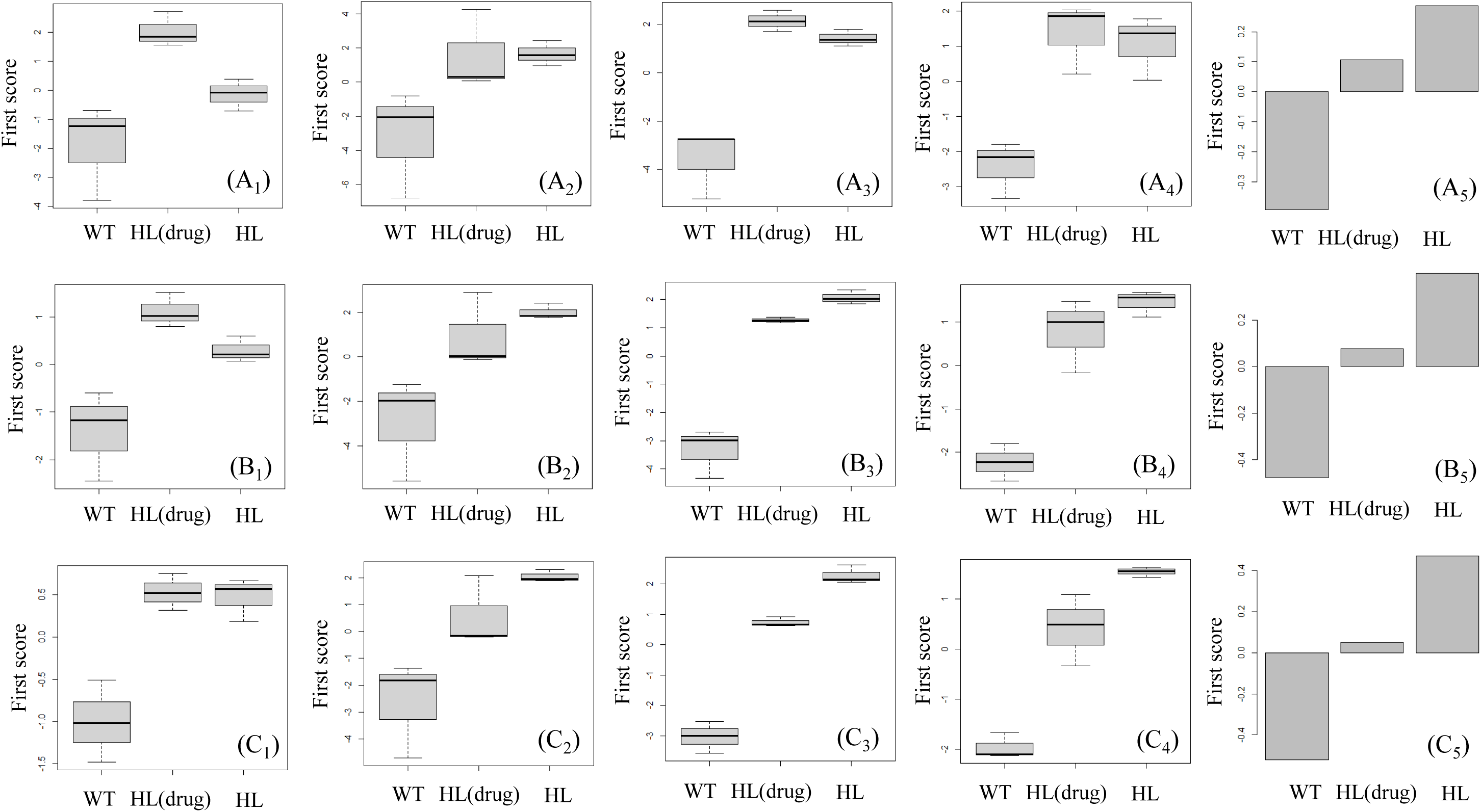
A box and whisker plot of first multiset PLS-ROG score for explanatory variables to wild type (WT), hyperlipidemia (HL) administered with drug and HL in (A_1_) plasma samples (τ=0.23), (A_2_) liver samples (τ=0.23), (A_3_) heart muscle samples (τ=0.23), (A_4_) brain samples (τ=0.23), in (B_1_) plasma samples (τ=0.24), (B_2_) liver samples (τ=0.24), (B_3_) heart muscle samples (τ=0.24), (B_4_) brain samples (τ=0.24), in (C_1_) plasma samples (τ=0.25), (C_2_) liver samples (τ=0.25), (C_3_) heart muscle samples (τ=0.25) and (C_4_) brain samples (τ=0.25). A bar plot of first multiset PLS-ROG score for response variable in (A_5_) τ=0.23, (B_5_) τ=0.24 and (C_5_) τ=0.25 to HC, mild and severe COVID-19 groups.

We have confirmed how the multiset PLS-ROG scores change with the parameter τ. Now, we focused on the results of multiset PLS-ROG in heart muscle samples because group difference and rank order of groups were more clear than other samples. To confirm how highly ranked metabolites correlated with the multiset PLS-ROG score were changed, we picked the top 3 ranked metabolites by multiset PLS-ROG loading at τ =0.23, 0.24 and 0.25 in heart muscle samples. The top 3 metabolites were carnosine (R=0.9489, p=9.476×10^-5^, q=0.0153) (Fig.9 (A)), citrulline (R=-0.9230, p=3.872×10^-4^, q=0.0232) (Fig.9 (B)) and O-acetylserine (R=0.9489, p=4.330×10^-4^, q=0.0232) (Fig.9 (C)) at τ=0.23, and citrulline (R=-0.8790, p=1.797×10^-3^, q=0.1202) (Fig.9 (A)), N6,N6,N6-trimethyllysine (R=-0.8626, p=2.760 × 10^-3^, q=0.1202) (Fig.9 (D)) and N,N-dimethylglycine (R=0.8503, p=3.681×10^-3^, q=0.1202) (Fig.9(E)) at τ=0.24 and N,N-dimethylglycine (R=0.9084, p=6.988×10^-4^, q=0.1125) (Fig.9 (E)), tyramine (R=0.8515, p=3.581 × 10^-3^, q=0.1987) (Fig.9 (F)) and Gly-Gly (R=-0.8385, p=4.744 × 10^-3^, q= 0.1987) (Fig.9 (G)) at τ=0.25.

**Figure 9.**
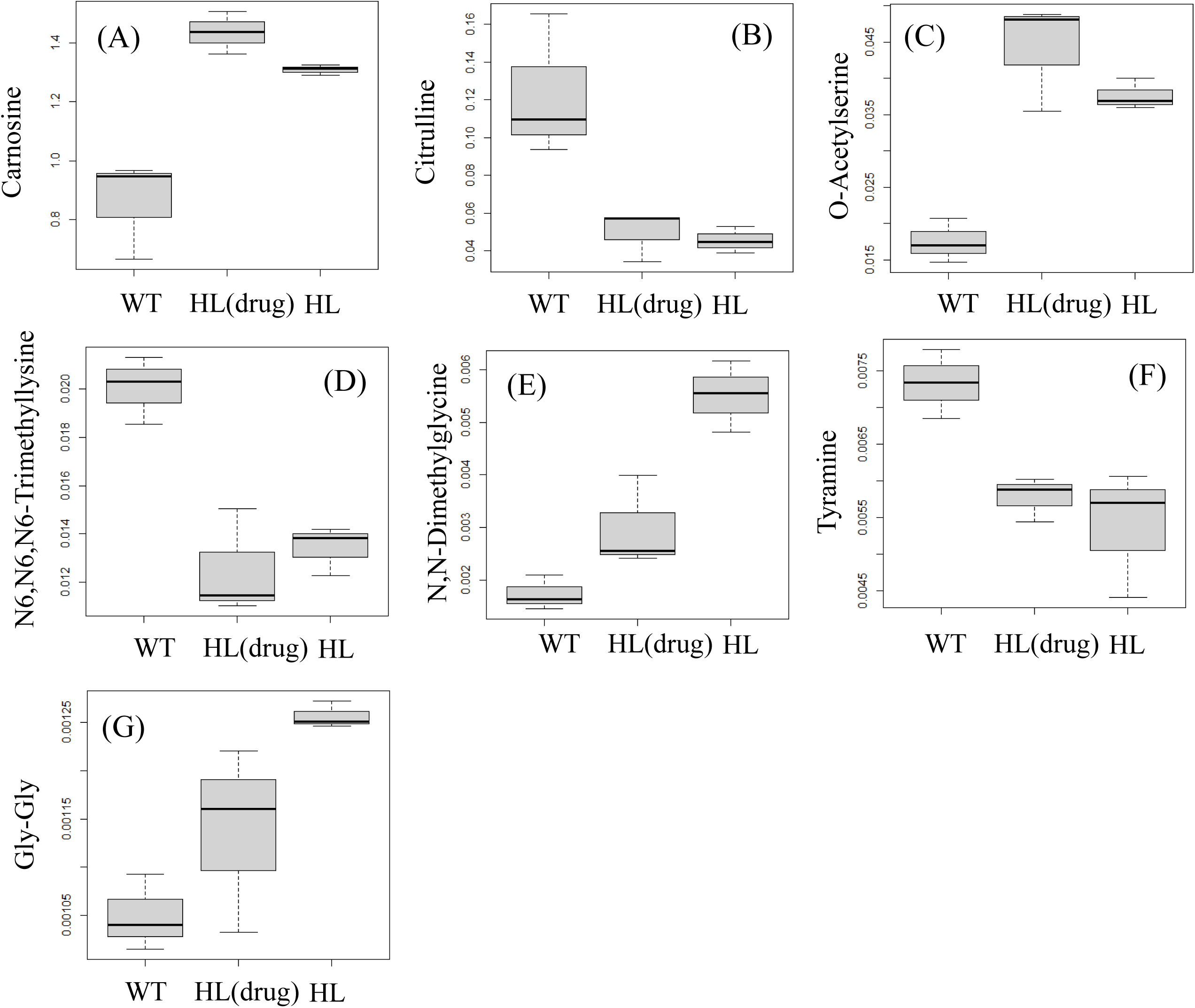
A box and whisker plot of (A) carnosine, (B) citrulline, (C) O-acetylserine, (D) N6,N6,N6-trimethyllysine, (E) N,N-dimethylglycine, (F) tyramine and (G) Gly-Gly to wild type (WT), hyperlipidemia (HL) administered with drug and HL in heart muscle samples.

We could not confirm metabolites that showed rank order of groups clearly at τ=0.23. Among them, group difference between wild type and other groups were shown clearly and rank order of groups were shown slightly in citrulline of second ranked metabolite (Fig9 (A)). At τ=0.24, citrulline was ranked first and N,N-dimethylglycine that showed rank order of groups clearly was ranked third. Furthermore, at τ=0.25, N,N-dimethylglycine was ranked first and Gly-Gly of third ranked metabolite also showed rank order of groups. And citrulline was forth ranked at τ=0.25. From these results, we could confirm that rank order of groups was clearer in highly ranked metabolites as τ is increased from 0.23 to 0.25.

About the significant metabolites selected by multiset PLS-ROG, dimethylglycine, Gly-Gly and citrulline were mentioned in the original paper. On the other hand, tyramine of second ranked metabolites at τ=0.25 that showed rank order of groups slightly was not mentioned. It was reported that tyramine was included in a variety of foods and was a compound that triggers hypertensive attacks [15].

## 4. Discussion

In this section, we would like to discuss some points to keep in mind for practical data analysis by using multiset PLS-ROG. For visualization of data, it is important to check both multiset PLS-ROG scores for explanatory variable and response variable because the pattern of multiset PLS-ROG scores for the explanatory and response variables sometimes differed each other. For example, in COVID-19 study, we could confirm that multiset PLS-ROG score for response variable showed severity of disease clearly, but that for explanatory variables did not show severity of disease (Fig.2). This becomes more important when we select significant compounds by using multiset PLS-ROG loading. In our previous study of PLS-ROG [10, 16], we could identify significant metabolites with rank order of groups by using PLS-ROG loading, if only the response variable showed rank order of the groups. On the other hands, we could identify compounds with rank order of groups by using multiset PLS-ROG loading only when both the explanatory and response variables showed rank order of groups. The reason we need to confirm both the explanatory and the response variables can be explained theoretically as follows, whereas PLS-ROG loading is defined as the correlation coefficient between PLS-ROG score for the response variable and each compound, multiset PLS-ROG loading is defined as the weighted sum of correlation coefficient between multiset PLS-ROG score of response variable and each compounds and correlation coefficient between multiset PLS-ROG score for other explanatory variable and each compounds.

We would like to discuss about the parameters of great impact in the multiset PLS-ROG to extract scores with rank order of groups. In our previous study of PLS-ROG, as the smoothing parameter κ increased, the rank order of groups in PLS-ROG score became clearer. On the other hand, in multiset PLS-ROG, not only the smoothing parameter κ increased but also the weight of connectivity τ between group information and each dataset increased, the correlation coefficient between multiset PLS-ROG score for response variable and each explanatory variable became higher. Consequently, we could confirm multiset PLS-ROG scores with rank order of groups clearly in this study of COVID-19 and hyperlipidemia. It was also commonly confirmed in COVID-19 and hyperlipidemia study that the multiset PLS-ROG scores swapped from the second multiset PLS-ROG score to the first multiset PLS-ROG score as the weight of connectivity between the groups and each dataset was increased. This also suggested that the effect of rank order of groups might become clearer in first multiset PLS-ROG score as the weight of the connectivity between group and each dataset increased.

Finally, we would like to discuss how to set the parameter for the weight of connection between each omics datasets or response variable τ. In COVID-19 study, multiset PLS-ROG score for explanatory variables at τ=0.23 was affected by common outlier samples but could not show rank order of groups. On the other hands, multiset PLS-ROG score for explanatory variables at τ=0.25 was not affected by outliers and showed rank order of groups. When we focused on only rank order of groups, the optimal parameter of τ might be 0.25. However, in the case of human clinical research such as COVID-19 study, it may be practical to consider the rank order of groups with some allowance for outliers. In that case, the optimal parameter τ might be 0.24 in which both the effect of outliers and the order of the groups. As well as COVID-19 study, we confirmed the effects of change for the parameter τ for the weight of connection between each metabolome data on multiset PLS-ROG score in hyperlipidemia study. In addition, there may be also considered various patterns of connection between datasets in hyperlipidemia study, for example, in cases where the heart muscle was connected to the liver but not to the plasma. Thus, there are more pattens of connection between group and each dataset not only how the weight of connection is set, but also connected or not. In practical data analysis, it would be important to subjectively determine the parameter τ for the weight of connection depending on the purpose. It will be necessary to further apply the multiset PLS-ROG to various data sets and further study how the parameter τ should be set.

## 5. Conclusion

We proposed multiset PLS-ROG to integrate multi-omics data with rank order of groups. We explored the effect of the parameters on the multiset PLS-ROG scores through practical data analysis of multi-omics data in COVID-19 study and multi-organ derived metabolome data in hyperlipidemia study. As a result, we found that both parameters of weight of connection τ between group and datasets and smoothing κ were important to extract scores with rank order of groups while only smoothing parameter is important in PLS-ROG. To identify compounds by statistical criteria, we clarified the statistical property of the weight vector in multiset PLS-ROG that was proportional to the weighted sum of correlation coefficient between multiset PLS-ROG score for response variable and each compound levels and multiset PLS-ROG score for explanatory variable and each compound levels. With this statistical property, we defined multiset PLS-ROG loading as the weighted correlation coefficient and could identify statistically significant compounds by using statistical hypothesis testing of multiset PLS-ROG loading.

## CRediT author statement

Hiroyuki Yamamoto: Conceptualization, Methodology, Software, Writing original draft and reviewing and Editing.

## Declaration of competing interest

The author is an employee of Human Metabolome Technologies, Inc.

## Supplementary

**Figure S1** First and second unsupervised multiset PLS scores in COVID-19 study. Three outlier samples of JBDZ2, JBDZ3 and JBDZ6 were blacked out. (A) A scatter plot of first and second unsupervised multiset PLS score for explanatory variables in proteome dataset. (◯) Healthy control (HC), (△) mild COVID-19, (□) severe COVID-19. (B) A scatter plot of first and second unsupervised multiset PLS score for explanatory variables in metabolome dataset.

**Figure S2** First and second multiset PLS scores (τ_1,2_=t_1,Y_=t_2,Y_=1/3) in COVID-19 study. Three outlier samples of JBDZ2, JBDZ3 and JBDZ6 were blacked out. (A_1_) A scatter plot of first and second multiset PLS score for explanatory variables in proteome dataset. (◯) Healthy control (HC), (△) mild COVID-19, (□) severe COVID-19. A box and whisker plot of (A_2_) first multiset PLS score and (A_3_) second multiset PLS score for explanatory variables to HC, mild and severe COVID-19 groups in proteome dataset. (B_1_) A scatter plot of first and second multiset PLS score for explanatory variables in metabolome dataset. A box and whisker plot of (B_2_) first multiset PLS score and (B_3_) second multiset PLS score for explanatory variables to HC, mild and severe COVID-19 groups in metabolome dataset. (C_1_) A scatter plot of first and second multiset PLS score for response variables. A bar plot of (C_2_) first multiset PLS score and (C_3_) second multiset PLS score for response variable to HC, mild and severe COVID-19 groups.

**Figure S3** First and second multiset PLS-ROG scores (τ_1,Y_=t_2,Y_=0.48) in COVID-19 study. Three outlier samples of JBDZ2, JBDZ3 and JBDZ6 were blacked out. (A_1_) A scatter plot of first and second multiset PLS-ROG score for explanatory variables in proteome dataset. (◯) Healthy control (HC), (△) mild COVID-19, (□) severe COVID-19. A box and whisker plot of (A_2_) first multiset PLS-ROG score and (A_3_) second multiset PLS-ROG score for explanatory variables to HC, mild and severe COVID-19 groups in proteome dataset. (B_1_) A scatter plot of first and second multiset PLS-ROG score for explanatory variables in metabolome dataset. A box and whisker plot of (B_2_) first multiset PLS-ROG score and (B_3_) second multiset PLS-ROG score for explanatory variables to HC, mild and severe COVID-19 groups in metabolome dataset. (C_1_) A scatter plot of first and second multiset PLS-ROG score for response variables. A bar plot of (C_2_) first multiset PLS-ROG score and (C_3_) second multiset PLS-ROG score for response variable to HC, mild and severe COVID-19 groups.

**Figure S4** First and second multiset PLS-ROG scores (τ_1,Y_=t_2,Y_=0.49) in COVID-19 study. Three outlier samples of JBDZ2, JBDZ3 and JBDZ6 were blacked out. (A_1_) A scatter plot of first and second multiset PLS-ROG score for explanatory variables in proteome dataset. (◯) Healthy control (HC), (△) mild COVID-19, (□) severe COVID-19. A box and whisker plot of (A_2_) first multiset PLS-ROG score and (A_3_) second multiset PLS-ROG score for explanatory variables to HC, mild and severe COVID-19 groups in proteome dataset. (B_1_) A scatter plot of first and second multiset PLS-ROG score for explanatory variables in metabolome dataset. A box and whisker plot of (B_2_) first multiset PLS-ROG score and (B_3_) second multiset PLS-ROG score for explanatory variables to HC, mild and severe COVID-19 groups in metabolome dataset. (C_1_) A scatter plot of first and second multiset PLS-ROG score for response variables. A bar plot of (C_2_) first multiset PLS-ROG score and (C_3_) second multiset PLS-ROG score for response variable to HC, mild and severe COVID-19 groups.

**Figure S5** First and second multiset PLS-ROG scores (τ_1,Y_=t_2,Y_=0.50) in COVID-19 study. Three outlier samples of JBDZ2, JBDZ3 and JBDZ6 were blacked out. (A_1_) A scatter plot of first and second multiset PLS-ROG score for explanatory variables in proteome dataset. (◯) Healthy control (HC), (△) mild COVID-19, (□) severe COVID-19. A box and whisker plot of (A_2_) first multiset PLS-ROG score and (A_3_) second multiset PLS-ROG score for explanatory variables to HC, mild and severe COVID-19 groups in proteome dataset. (B_1_) A scatter plot of first and second multiset PLS-ROG score for explanatory variables in metabolome dataset. A box and whisker plot of (B_2_) first multiset PLS-ROG score and (B_3_) second multiset PLS-ROG score for explanatory variables to HC, mild and severe COVID-19 groups in metabolome dataset. (C_1_) A scatter plot of first and second multiset PLS-ROG score for response variables. A bar plot of (C_2_) first multiset PLS-ROG score and (C_3_) second multiset PLS-ROG score for response variable to HC, mild and severe COVID-19 groups.

**Figure S6** A box and whisker plot of 5-methyluridine to HC, mild and severe COVID-19 groups. **Figure S7** First and second multiset PLS-ROG scores (τ = 0.21) in hyperlipidemia study. (◯) Wild type (WT), (△) hyperlipidemia (HL) rabbit administered with drug, (□) HL rabbit. (A_1_) A scatter plot of first and second multiset PLS-ROG score for explanatory variables in plasma samples. A box and whisker plot of (A_2_) first multiset PLS-ROG score and (A_3_) second multiset PLS-ROG score for explanatory variables to WT, HL administered with drug and HL in plasma samples. (B_1_) A scatter plot of first and second multiset PLS-ROG score for explanatory variables to WT, HL administered with drug and HL in liver samples. A box and whisker plot of (B_2_) first multiset PLS-ROG score (B_3_) second multiset PLS-ROG score for explanatory variables to WT, HL administered with drug and HL in liver samples. (C_1_) A scatter plot of first and second multiset PLS-ROG score for explanatory variables in heart muscle samples. A box and whisker plot of (C_2_) first multiset PLS-ROG score and (C_3_) second multiset PLS-ROG score for explanatory variables to WT, HL administered with drug and HL in heart muscle samples. (D_1_) A scatter plot of first and second multiset PLS-ROG score for explanatory variables in brain samples. A box and whisker plot of (D_2_) first multiset PLS-ROG score and (D_3_) second multiset PLS-ROG score for explanatory variables to WT, HL administered with drug and HL in brain samples. (E_1_) A scatter plot of first and second multiset PLS-ROG score for response variables. A bar plot of (E_2_) first multiset PLS-ROG score and (E_3_) second multiset PLS-ROG score for response variable to WT, HL administered with drug and HL.

**Figure S8** First and second multiset PLS-ROG scores (τ = 0.22) in hyperlipidemia study. (◯) Wild type (WT), (△) hyperlipidemia (HL) rabbit administered with drug, (□) HL rabbit. (A_1_) A scatter plot of first and second multiset PLS-ROG score for explanatory variables in plasma samples. A box and whisker plot of (A_2_) first multiset PLS-ROG score and (A_3_) second multiset PLS-ROG score for explanatory variables to WT, HL administered with drug and HL in plasma samples. (B_1_) A scatter plot of first and second multiset PLS-ROG score for explanatory variables to WT, HL administered with drug and HL in liver samples. A box and whisker plot of (B_2_) first multiset PLS-ROG score (B_3_) second multiset PLS-ROG score for explanatory variables to WT, HL administered with drug and HL in liver samples. (C_1_) A scatter plot of first and second multiset PLS-ROG score for explanatory variables in heart muscle samples. A box and whisker plot of (C_2_) first multiset PLS-ROG score and (C_3_) second multiset PLS-ROG score for explanatory variables to WT, HL administered with drug and HL in heart muscle samples. (D_1_) A scatter plot of first and second multiset PLS-ROG score for explanatory variables in brain samples. A box and whisker plot of (D_2_) first multiset PLS-ROG score and (D_3_) second multiset PLS-ROG score for explanatory variables to WT, HL administered with drug and HL in brain samples. (E_1_) A scatter plot of first and second multiset PLS-ROG score for response variables. A bar plot of (E_2_) first multiset PLS-ROG score and (E_3_) second multiset PLS-ROG score for response variable to WT, HL administered with drug and HL.

**Figure S9** First and second multiset PLS-ROG scores (τ = 0.23) in hyperlipidemia study. (◯) Wild type (WT), (△) hyperlipidemia (HL) rabbit administered with drug, (□) HL rabbit. (A_1_) A scatter plot of first and second multiset PLS-ROG score for explanatory variables in plasma samples. A box and whisker plot of (A_2_) first multiset PLS-ROG score and (A_3_) second multiset PLS-ROG score for explanatory variables to WT, HL administered with drug and HL in plasma samples. (B_1_) A scatter plot of first and second multiset PLS-ROG score for explanatory variables to WT, HL administered with drug and HL in liver samples. A box and whisker plot of (B_2_) first multiset PLS-ROG score (B_3_) second multiset PLS-ROG score for explanatory variables to WT, HL administered with drug and HL in liver samples. (C_1_) A scatter plot of first and second multiset PLS-ROG score for explanatory variables in heart muscle samples. A box and whisker plot of (C_2_) first multiset PLS-ROG score and (C_3_) second multiset PLS-ROG score for explanatory variables to WT, HL administered with drug and HL in heart muscle samples. (D_1_) A scatter plot of first and second multiset PLS-ROG score for explanatory variables in brain samples. A box and whisker plot of (D_2_) first multiset PLS-ROG score and (D_3_) second multiset PLS-ROG score for explanatory variables to WT, HL administered with drug and HL in brain samples. (E_1_) A scatter plot of first and second multiset PLS-ROG score for response variables. A bar plot of (E_2_) first multiset PLS-ROG score and (E_3_) second multiset PLS-ROG score for response variable to WT, HL administered with drug and HL.

**Figure S10** First and second multiset PLS-ROG scores (τ = 0.24) in hyperlipidemia study. (◯) Wild type (WT), (△) hyperlipidemia (HL) rabbit administered with drug, (□) HL rabbit. (A_1_) A scatter plot of first and second multiset PLS-ROG score for explanatory variables in plasma samples. A box and whisker plot of (A_2_) first multiset PLS-ROG score and (A_3_) second multiset PLS-ROG score for explanatory variables to WT, HL administered with drug and HL in plasma samples. (B_1_) A scatter plot of first and second multiset PLS-ROG score for explanatory variables to WT, HL administered with drug and HL in liver samples. A box and whisker plot of (B_2_) first multiset PLS-ROG score (B_3_) second multiset PLS-ROG score for explanatory variables to WT, HL administered with drug and HL in liver samples. (C_1_) A scatter plot of first and second multiset PLS-ROG score for explanatory variables in heart muscle samples. A box and whisker plot of (C_2_) first multiset PLS-ROG score and (C_3_) second multiset PLS-ROG score for explanatory variables to WT, HL administered with drug and HL in heart muscle samples. (D_1_) A scatter plot of first and second multiset PLS-ROG score for explanatory variables in brain samples. A box and whisker plot of (D_2_) first multiset PLS-ROG score and (D_3_) second multiset PLS-ROG score for explanatory variables to WT, HL administered with drug and HL in brain samples. (E_1_) A scatter plot of first and second multiset PLS-ROG score for response variables. A bar plot of (E_2_) first multiset PLS-ROG score and (E_3_) second multiset PLS-ROG score for response variable to WT, HL administered with drug and HL.

**Figure S11** First and second multiset PLS-ROG scores (τ = 0.25) in hyperlipidemia study. (◯) Wild type (WT), (△) hyperlipidemia (HL) rabbit administered with drug, (□) HL rabbit. (A_1_) A scatter plot of first and second multiset PLS-ROG score for explanatory variables in plasma samples. A box and whisker plot of (A_2_) first multiset PLS-ROG score and (A_3_) second multiset PLS-ROG score for explanatory variables to WT, HL administered with drug and HL in plasma samples. (B_1_) A scatter plot of first and second multiset PLS-ROG score for explanatory variables to WT, HL administered with drug and HL in liver samples. A box and whisker plot of (B_2_) first multiset PLS-ROG score (B_3_) second multiset PLS-ROG score for explanatory variables to WT, HL administered with drug and HL in liver samples. (C_1_) A scatter plot of first and second multiset PLS-ROG score for explanatory variables in heart muscle samples. A box and whisker plot of (C_2_) first multiset PLS-ROG score and (C_3_) second multiset PLS-ROG score of explanatory variables to WT, HL administered with drug and HL in heart muscle samples. (D_1_) A scatter plot of first and second multiset PLS-ROG score for explanatory variables in brain samples. A box and whisker plot of (D_2_) first multiset PLS-ROG score and (D_3_) second multiset PLS-ROG score for explanatory variables to WT, HL administered with drug and HL in brain samples. (E_1_) A scatter plot of first and second multiset PLS-ROG score for response variables. A bar plot of (E_2_) first multiset PLS-ROG score and (E_3_) second multiset PLS-ROG score for response variable to WT, HL administered with drug and HL.

## Notes

### Summary of Updates

We have revised the definition of the loadings for the multiset PLS-ROG. The results of the analysis have been revised accordingly.

## References

[1] S. Huang, K. Chaudhary, L.X. Garmire, More is better: Recent progress in multi-omics data integration methods, Front. Genet. 8 (2017) 1–12. 10.3389/fgene.2017.00084.

[2] F.R. Pinu, D.J. Beale, A.M. Paten, K. Kouremenos, S. Swarup, H.J. Schirra, D. Wishart, Systems biology and multi-omics integration: Viewpoints from the metabolomics research community, Metabolites. 9 (2019) 1–31. 10.3390/metabo9040076.

[3] N. Ishii, K. Nakahigashi, T. Baba, M. Robert, T. Soga, A. Kanai, T. Hirasawa, M. Naba, K. Hirai, A. Hoque, P.Y. Ho, Y. Kakazu, K. Sugawara, S. Igarashi, S. Harada, T. Masuda, N. Sugiyama, T. Togashi, M. Hasegawa, Y. Takai, K. Yugi, K. Arakawa, N. Iwata, Y. Toya, Y. Nakayama, T. Nishioka, K. Shimizu, H. Mori, M. Tomita, Multiple high-throughput analyses monitor the response of E. coli to perturbations, Science (80-.). 316 (2007) 593–597. 10.1126/science.1132067.

[4] K. Yugi, H. Kubota, A. Hatano, S. Kuroda, Trans-Omics: How To Reconstruct Biochemical Networks Across Multiple “Omic” Layers, Trends Biotechnol. 34 (2016) 276–290. 10.1016/j.tibtech.2015.12.013.

[5] J. Lloyd-Price, C. Arze, A.N. Ananthakrishnan, M. Schirmer, J. Avila-Pacheco, T.W. Poon, E. Andrews, N.J. Ajami, K.S. Bonham, C.J. Brislawn, D. Casero, H. Courtney, A. Gonzalez, T.G. Graeber, A.B. Hall, K. Lake, C.J. Landers, H. Mallick, D.R. Plichta, M. Prasad, G. Rahnavard, J. Sauk, D. Shungin, Y. Vázquez-Baeza, R.A. White, J. Bishai, K. Bullock, A. Deik, C. Dennis, J.L. Kaplan, H. Khalili, L.J. McIver, C.J. Moran, L. Nguyen, K.A. Pierce, R. Schwager, A. Sirota-Madi, B.W. Stevens, W. Tan, J.J. ten Hoeve, G. Weingart, R.G. Wilson, V. Yajnik, J. Braun, L.A. Denson, J.K. Jansson, R. Knight, S. Kugathasan, D.P.B. McGovern, J.F. Petrosino, T.S. Stappenbeck, H.S. Winter, C.B. Clish, E.A. Franzosa, H. Vlamakis, R.J. Xavier, C. Huttenhower, Multi-omics of the gut microbial ecosystem in inflammatory bowel diseases, Nature. 569 (2019) 655–662. 10.1038/s41586-019-1237-9.

[6] Y. Hasin, M. Seldin, A. Lusis, Multi-omics approaches to disease, Genome Biol. 18 (2017) 1–15. 10.1186/s13059-017-1215-1.

7. [7] A. Csala, A.H. Zwinderman, “Multivariate Statistical Methods for High-Dimensional Multiset Omics Data Analysis”, Computational Biology, Husi H, editor., Brisbane (AU), Codon Publications (2019). 10.15586/computationalbiology.2019.ch5

[8] F. Rohart, B. Gautier, A. Singh, K.A. Lê Cao, mixOmics: An R package for ‘omics feature selection and multiple data integration, PLoS Comput. Biol. 13 (2017) 1–19. 10.1371/journal.pcbi.1005752.

[9] H. Yamamoto, T. Fujimori, H. Sato, G. Ishikawa, K. Kami, Y. Ohashi, Statistical hypothesis testing of factor loading in principal component analysis and its application to metabolite set enrichment analysis, BMC Bioinformatics. 15 (2014). 10.1186/1471-2105-15-51.

[10] H. Yamamoto, PLS-ROG: Partial least squares with rank order of groups, J. Chemom. 31 (2017) e2883. 10.1002/cem.2883.

[11] H. Yamamoto, PLS-ROG: Partial least squares with rank order of groups, J. Chemom. 31 (2017) e2883. 10.1002/cem.2883.

[12] B. Shen, X. Yi, Y. Sun, X. Bi, J. Du, C. Zhang, S. Quan, F. Zhang, R. Sun, L. Qian, W. Ge, W. Liu, S. Liang, H. Chen, Y. Zhang, J. Li, J. Xu, Z. He, B. Chen, J. Wang, H. Yan, Y. Zheng, D. Wang, J. Zhu, Z. Kong, Z. Kang, X. Liang, X. Ding, G. Ruan, N. Xiang, X. Cai, H. Gao, L. Li, S. Li, Q. Xiao, T. Lu, Y. Zhu, H. Liu, H. Chen, T. Guo, Proteomic and Metabolomic Characterization of COVID-19 Patient Sera, Cell. 182 (2020) 59–72.e15. 10.1016/j.cell.2020.05.032.

[13] T. Ooga, H. Sato, A. Nagashima, K. Sasaki, M. Tomita, T. Soga, Y. Ohashi, Metabolomic anatomy of an animal model revealing homeostatic imbalances in dyslipidaemia, Mol. Biosyst. 7 (2011) 1217–1223. 10.1039/c0mb00141d.

[14] P. Amarenco, J. Bogousslavsky, A. 3rd Callahan, L.B. Goldstein, M. Hennerici, A.E. Rudolph, H. Sillesen, L. Simunovic, M. Szarek, K.M.A. Welch, J.A. Zivin, High-dose atorvastatin after stroke or transient ischemic attack., N. Engl. J. Med. 355 (2006) 549–559. 10.1056/NEJMoa061894.

[15] S.L. Rice, R.R. Eitenmiller, P.E. Koehler, Biologically Active Amines in Food: A Review, J. Milk Food Technol. 39 (1976) 353–358. 10.4315/0022-2747-39.5.353

[16] E. Sasaki, H. Yamamoto, T. Asari, R. Matsuta, S. Ota, Y. Kimura, S. Sasaki, K. Ishibashi, Y. Yamamoto, K. Kami, M. Ando, E. Tsuda, Y. Ishibashi, Metabolomics with severity of radiographic knee osteoarthritis and early phase synovitis in middle-aged women from the Iwaki Health Promotion Project: a cross-sectional study, Arthritis Res. Ther. 24 (2022) 1–11. 10.1186/s13075-022-02830-w.

