## Supplementary Figures for "Multiset partial least squares with rank order of groups for integrating multi-omics data": FIGS3.pptx

### Slide 1
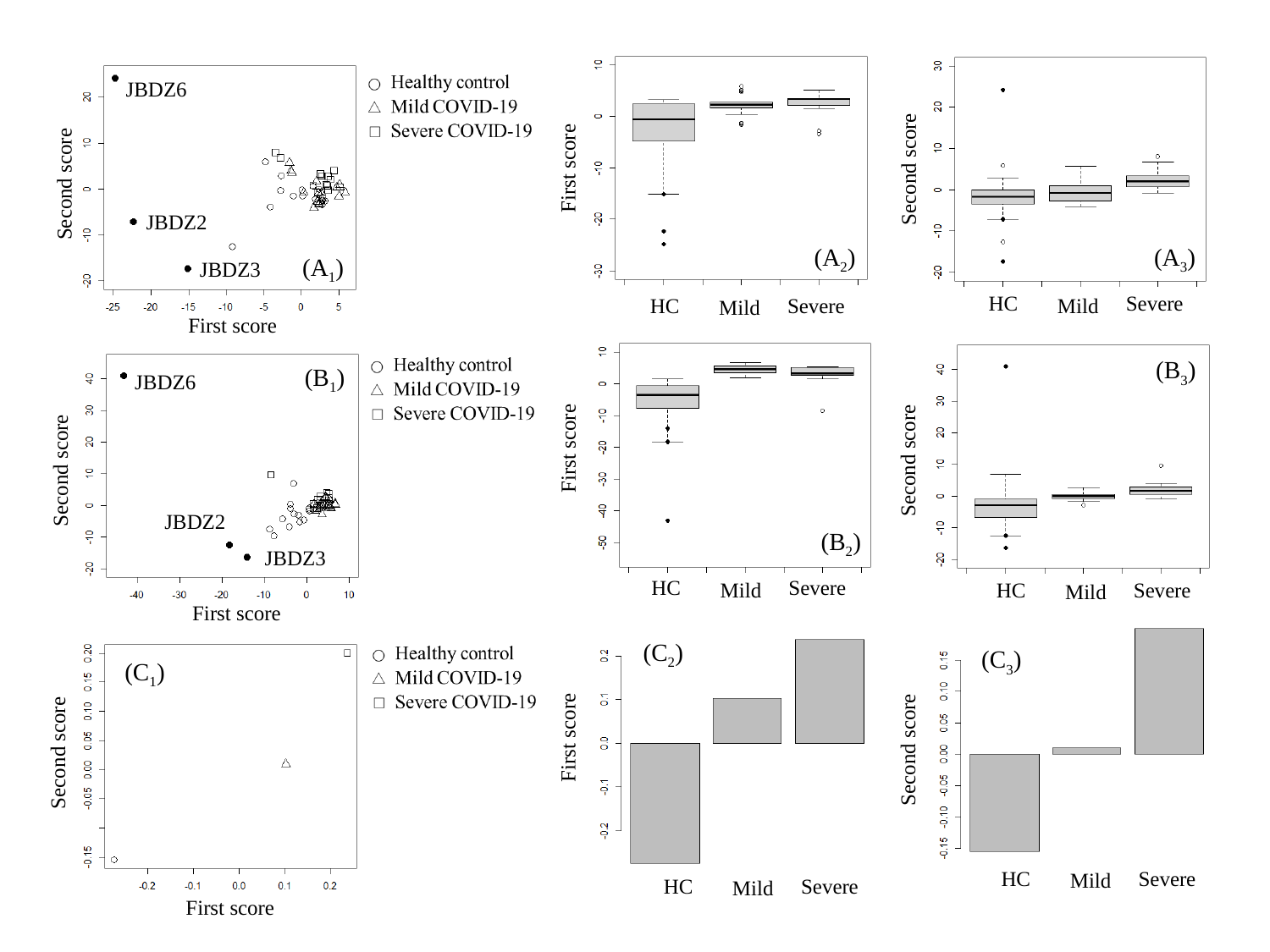

JBDZ6
First score
Second score
Second score
JBDZ2
(A_2_)
(A_3_)
(A_1_)
JBDZ3
HC
Severe
Mild
HC
Severe
Mild
First score
(B_3_)
(B_1_)
JBDZ6
First score
Second score
Second score
JBDZ2
(B_2_)
JBDZ3
HC
Severe
Mild
HC
Severe
Mild
First score
(C_2_)
(C_3_)
(C_1_)
First score
Second score
Second score
HC
Severe
Mild
HC
Severe
Mild
First score
