## Supplementary Figures for "Multiset partial least squares with rank order of groups for integrating multi-omics data": FIGS4.pptx

### Slide 1
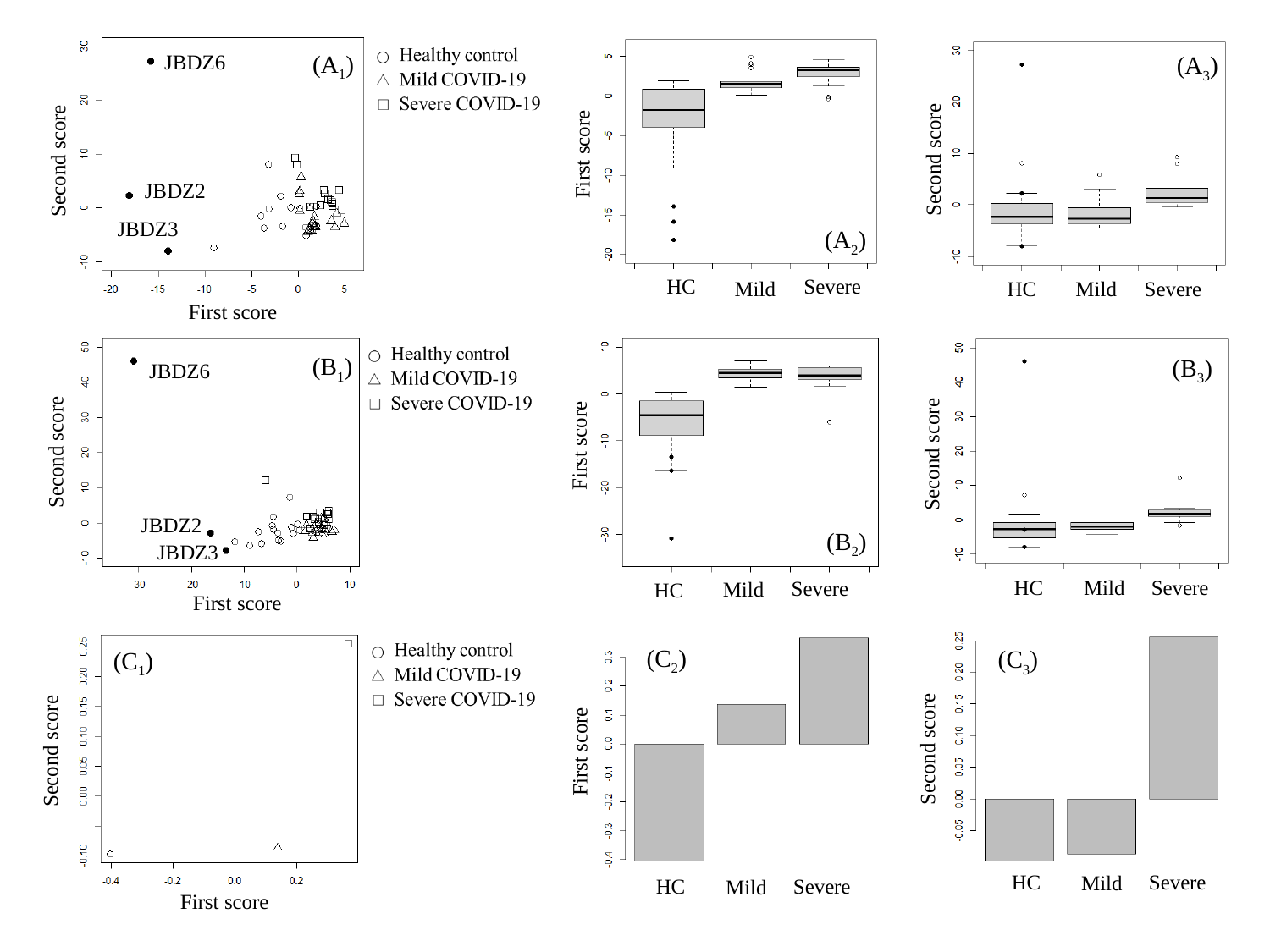

JBDZ6
(A_1_)
(A_3_)
First score
Second score
Second score
JBDZ2
JBDZ3
(A_2_)
HC
Severe
Mild
Mild
HC
Severe
First score
(B_1_)
(B_3_)
JBDZ6
First score
Second score
Second score
JBDZ2
(B_2_)
JBDZ3
Mild
HC
Severe
Severe
Mild
HC
First score
(C_2_)
(C_3_)
(C_1_)
Second score
Second score
First score
HC
Severe
Mild
HC
Severe
Mild
First score
