## Supplementary Figures for "Multiset partial least squares with rank order of groups for integrating multi-omics data": FIGS11.pptx

### Slide 1
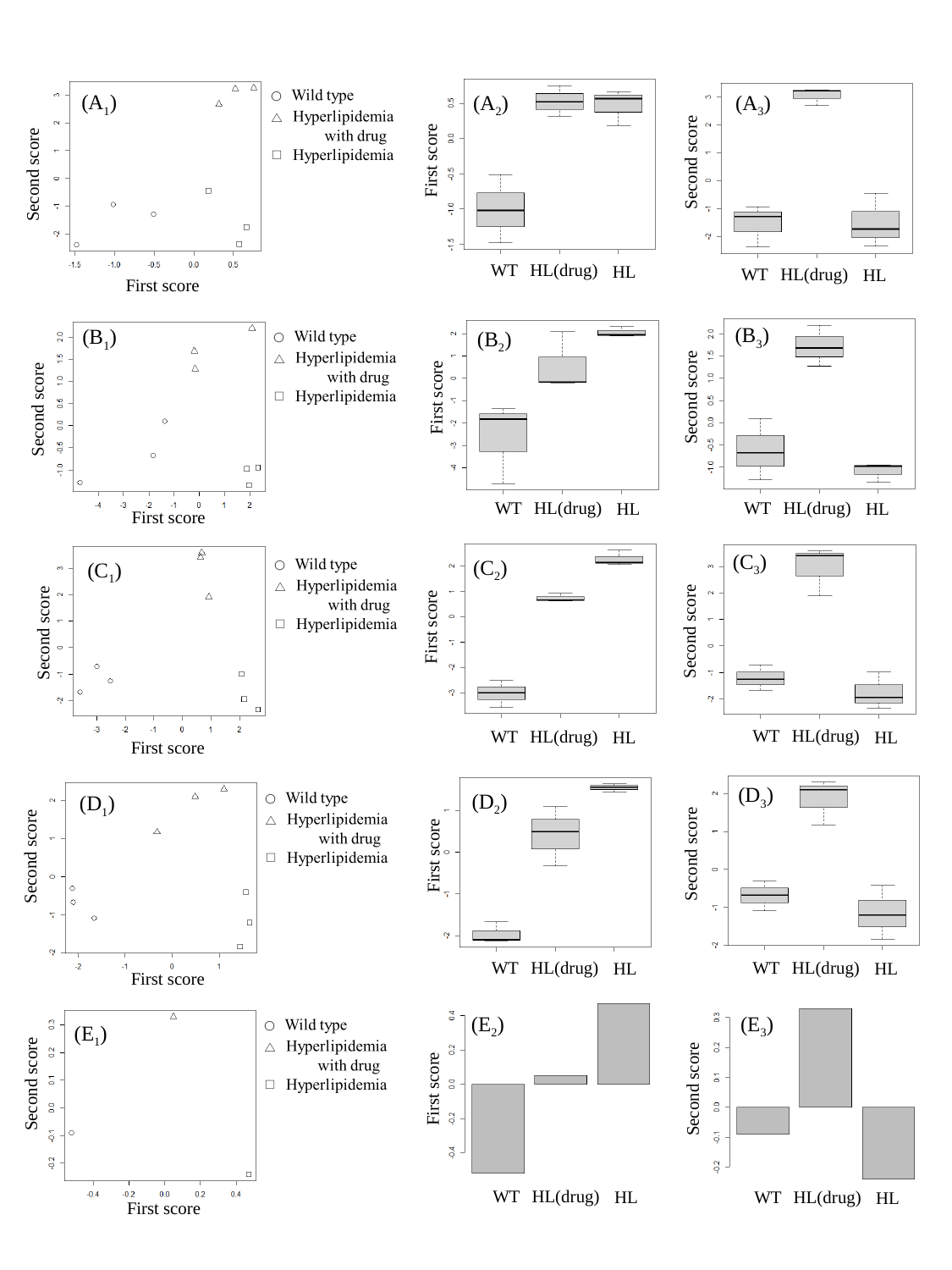

(A_1_)
(A_2_)
(A_3_)
First score
Second score
Second score
WT
HL(drug)
HL
WT
HL(drug)
HL
First score
(B_3_)
(B_1_)
(B_2_)
First score
Second score
Second score
WT
HL(drug)
WT
HL(drug)
HL
HL
First score
(C_3_)
(C_2_)
(C_1_)
First score
Second score
Second score
WT
HL(drug)
WT
HL(drug)
HL
HL
First score
(D_3_)
(D_2_)
(D_1_)
Second score
First score
Second score
WT
HL(drug)
WT
HL(drug)
HL
HL
First score
(E_2_)
(E_3_)
(E_1_)
Second score
Second score
First score
WT
HL(drug)
WT
HL(drug)
HL
HL
First score
