## Supplementary figures and images for "Multiset partial least squares with rank order of groups for integrating multi-omics data"

### FIGS1.pptx

#### Slide 1
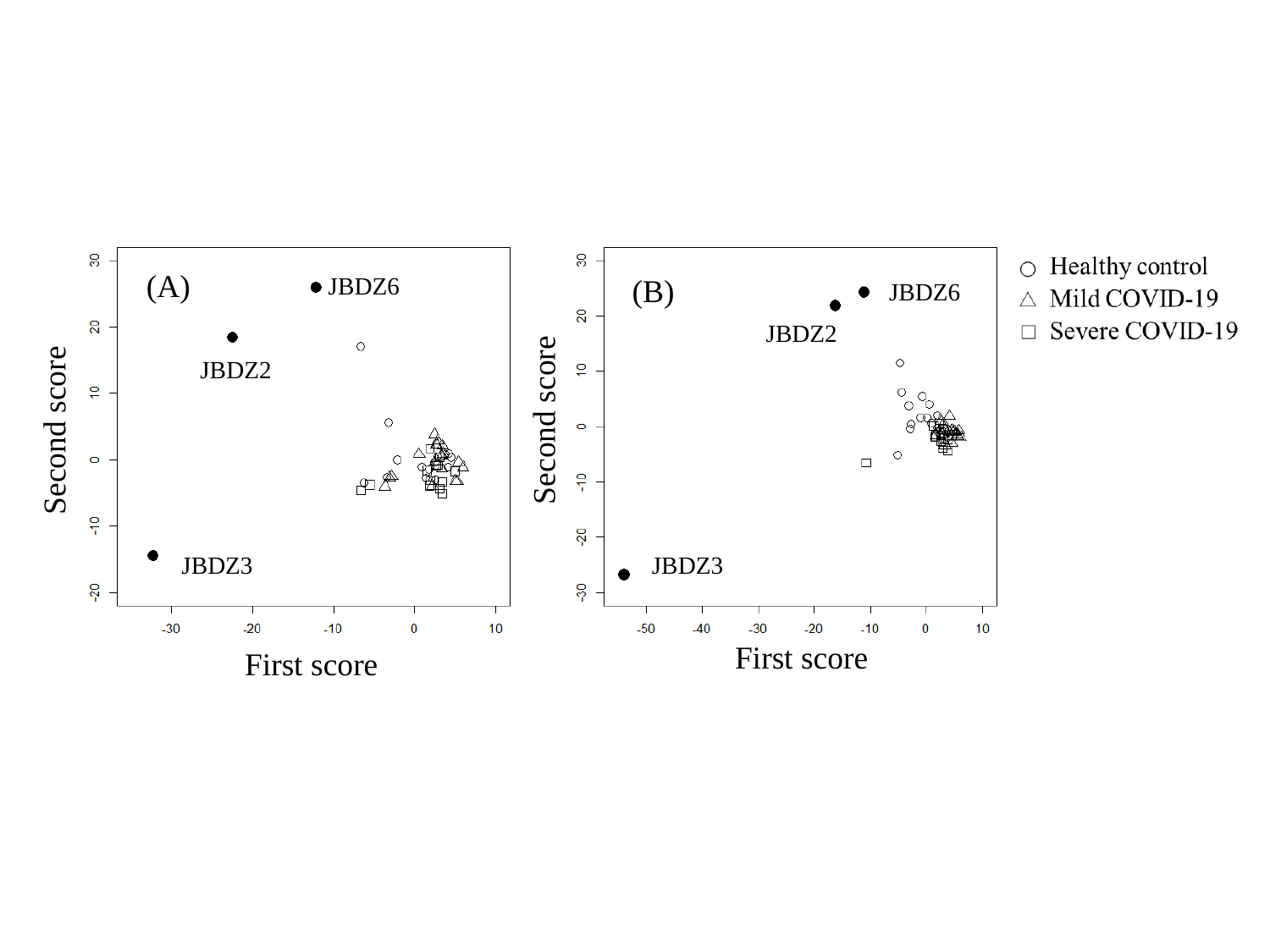

(A)
(B)
JBDZ6
JBDZ6
JBDZ2
JBDZ2
Second score
Second score
JBDZ3
JBDZ3
First score
First score

### FIGS6.pptx

#### Slide 1
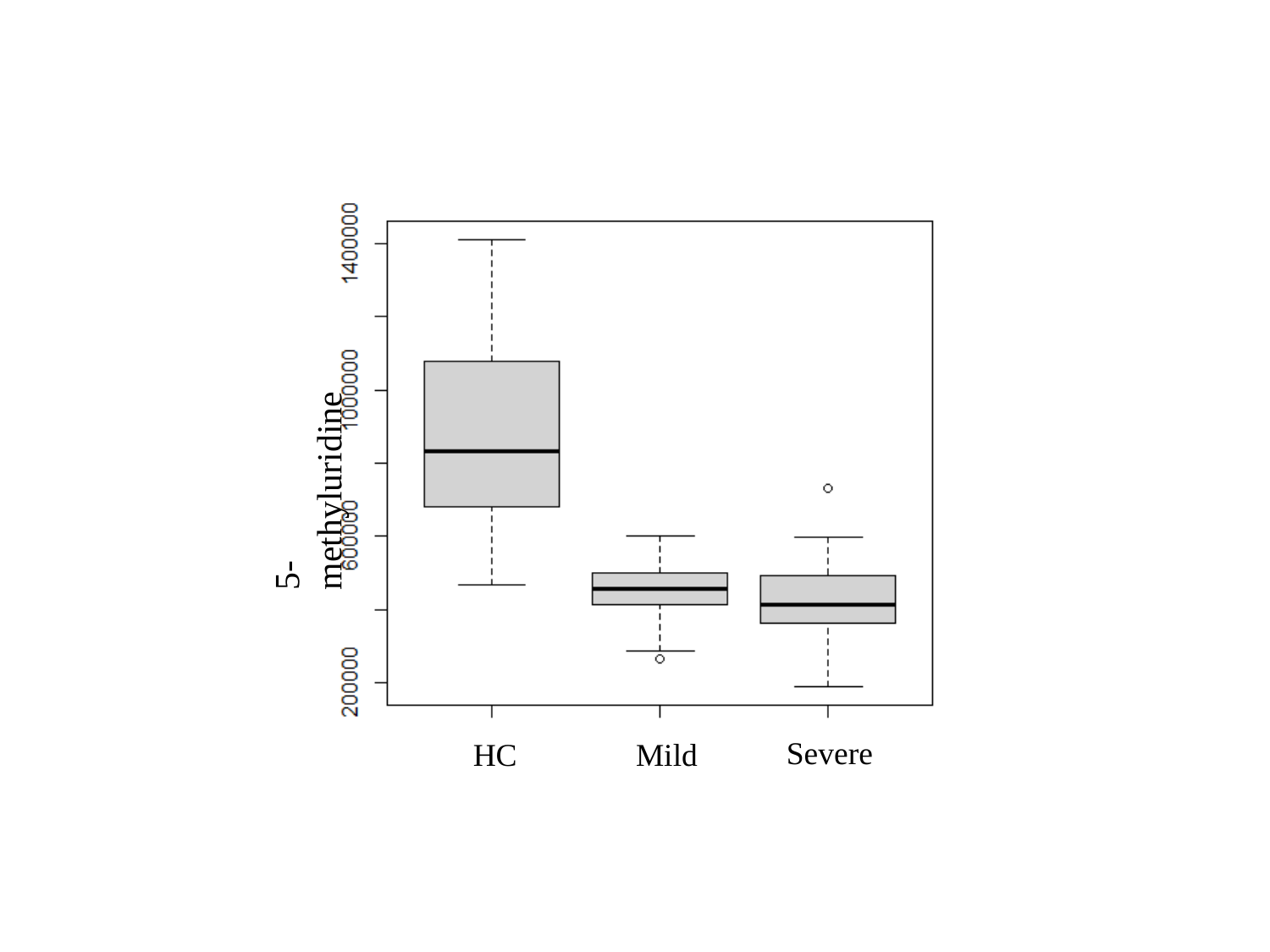

5-methyluridine
Severe
Mild
HC
